# Origin and transport of bacterial hopanoids

**DOI:** 10.1101/2025.07.29.667474

**Authors:** Evan C. Lawrence, Huiqiao Pan, Noah Ollikainen, Brittany J. Belin

## Abstract

Hopanoids are a class of steroid-like lipids that fortify bacterial membranes. Derivatives of hopanoids known as “geo-hopanes” are abundant in sediments and are signatures of ancient bacteria, yet there are conflicting views on their use as markers of specific taxa or environments. Here we analyze conservation of hopanoid biosynthesis across bacterial genomes using modern taxonomic tools. We find that hopanoids most likely originated in an ancestor of marine alphaproteobacteria, and that tolerance of high osmolarity is the most common feature of hopanoid-producing strains. Additionally, microsynteny with hopanoid-related loci revealed new hopanoid-associated gene families, which we term *hpnT* and *hpnS*. Structural predictions and the restriction of these gene families to diderms suggests they participate in hopanoid trafficking, potentially forming a pathway analogous to Mla proteins in *E. coli*.

## INTRODUCTION

Hopanoids are steroid-like isoprenoid lipids produced by bacteria^1,2^. In sediments, ancient hopanoid derivatives (“geo-hopanes”) are abundant, with the oldest biomarkers dating to >1.6 Ga^3^. Their longevity in geologic time is due to their highly stable, pentacyclic hydrocarbon core, generated by consolidation and cyclization of six 5-carbon isoprene units. The simplest hopanoids contain only this 30-carbon core, sometimes with minor modifications (methylation or C-C bond desaturation), and are known as the “short” or C_30_ class. Hopanoids with ribose-derived side chains form the “extended” or C_35_ class. The C_35_ hopanoids are structurally diverse, including the addition of hydroxyl or amine moieties or covalent attachment to other biomolecules.

Many studies have examined the extent to which hopanoid biomarkers can be used to infer the identities of ancient microbes and their environments (reviewed in ^4^). Such inferences rely on understanding which bacteria produce which hopanoids, with the assumption that the distribution of biological processes across phyla – and their compatibilities with certain environments (*e.g.*, oxic vs. anoxic habitats) – are conserved throughout time. Early *in vivo* studies hypothesized that geo-hopanes are markers of the advent of oxygenic photosynthesis. Specifically, hopanoids methylated at the C2 position (2Me-hopanoids) by the enzyme HpnP have been used as markers for marine cyanobacteria^3^, and the proliferation of 2Me-hopanoid derivatives during oceanic anoxic events (OAEs) led to the suggestion that marine cyanobacteria expanded in these periods^5^.

Increased availability of bacterial genomes and metagenomes, and the discovery of hopanoid biosynthesis enzymes (**Table 1**), have broadened our understanding of where and by which species hopanoids can be produced^6,7^. We now know that hopanoid-producing organisms are taxonomically diverse, making it difficult to classify geo-hopanes as markers of specific metabolisms or environmental contexts. The degree to which hopanoids can be used as markers of marine cyanobacteria remains an active subject of debate. Some genomics-based studies argue against use of hopanoids as cyanobacterial markers, suggesting that 2-Me hopanoids originated in the alphaproteobacteria^8^ and that hopanoids are more generally markers for nitrogen cycling^7,9–12^. Other works have reinforced the connection between hopanoids and cyanobacteria ^13^. Most recently it was suggested that 2-Me hopanoids can be used as cyanobacterial markers prior to 750 Ma^14^, based on the authors’ conclusions that (i) the evolutionary relationship between cyanobacterial hopanoid C2-methylases (*hpnP*) closely matches that of cyanobacteria genomes, indicating that *hpnP* was present in the last common cyanobacterial ancestor, and (ii) *hpnP*-encoding proteobacteria evolved much later than *hpnP*-encoding cyanobacteria^15^.

**Table 1.** Hopanoid-associated protein domain profiles in the Pfam/InterPro, TIGRFam, Kegg Ontology, and MetaCyc databases. *IPR005814 contains other aminotransferases and is not specific for HpnO/HpnQ. In the TIGRFAM database, an entry under hopanoid lipid biosynthesis pathway (GenProp0761) named HpnO is annotated with “VacJ-like lipoprotein”/PF04333. These terms do not refer to the same protein; use of HpnO/HpnQ to refer to the enzyme responsible for aminobacteriohopanetetrol production is the more common designation. The HpnO protein associated with GenProp0761 in TIGRFAM is likely the protein we name HpnS in this manuscript.

| Protein | Function | Pfam / Interpro | TIGRFam | Kegg Ontology |
| --- | --- | --- | --- | --- |
| SHC (HpnF) | C<sub>30</sub> hopanoid production<sup>69</sup> | pfam13249, pfam13243, IPR006400 | TIGR01507 | KO:K06045 |
| HpnG | production of C<sub>35</sub> hopanoid ribosylhopane<sup>70,71</sup> | IPR017831 | TIGR03468 | - |
| HpnH | production of C<sub>35</sub> hopanoid adenosylhopane<sup>70,72</sup> | IPR017833 | TIGR03470 | - |
| HpnI | production of C<sub>35</sub> hopanoid N-acetylglucosaminyl bacteriohopanetetrol (BHT)<sup>73</sup> | IPR017835 | TIGR03472 | - |
| HpnJ | production of C<sub>35</sub> hopanoid BHT cyclitol ether<sup>73</sup> | IPR017834 | TIGR03471 | - |
| HpnK | production of C<sub>35</sub> hopanoid glucosaminyl BHT<sup>73</sup> | IPR017836 | TIGR03473 | - |
| HpnL | Unknown | - | TIGR03476 | - |
| HpnM | Unknown | IPR017842 | TIGR03481 | - |
| HpnN | Inner membrane hopanoid transport<sup>74,75</sup> | IPR017841 | TIGR03480 | - |
| HpnO/<br>HpnQ | production of C<sub>35</sub> hopanoid aminobacteriohopanetetrol<sup>70</sup> | IPR005814* | “HpnO” under GenProp0761* |  |
| HpnP | C-2 methylation of hopanoids<sup>76</sup> | IPR034530 | - | KO:K22705 |
| HpnR | C-3 methylation of hopanoids<sup>77</sup> | IPR027564 | TIGR04367 | KO:K22704 |

Here we analyze the distribution of the hopanoid biosynthesis enzymes SHC, HpnH, HpnP, and HpnR across sequenced bacteria, incorporating an updated understanding of bacterial taxonomic relationships and comprehensive experimental data. Our results suggest that the capacity to produce hopanoids likely originated in an ancestor of marine alphaproteobacteria, and that tolerance of high osmolarity environments, nitrogen cycling, and single-carbon metabolism are common to hopanoid-producing species. Additionally, by evaluating microsynteny between known hopanoid-related genes and uncharacterized bacterial genes, we identify two new hopanoid-associated families, which we term *hpnT* and *hpnS*. Structural predictions and the restriction of these gene families to diderms suggests they participate in hopanoid trafficking, potentially forming a pathway analogous to Mla proteins in *E. coli* and other species.

## RESULTS

### Benchmarking annotation approaches for hopanoid-associated genes

Determining the distribution of hopanoids across bacteria requires a robust approach for annotation of hopanoid biosynthesis genes. Prevailing methods for gene annotation include comparison to reference gene sets and, for protein-coding genes, identification of conserved protein domains. The latter approach relies on profile hidden Markov models (HMM), and there are multiple domain profiles for most known hopanoid biosynthesis proteins across the InterPro, Pfam, TIGRFAM, and Kegg Ontology databases (**Table 1**). Domain assignments based on these profile HMMs are not manually curated and have not been benchmarked against experimental data.

To evaluate the accuracy of each hopanoid-associated profile HMM, we assembled a set of experimentally validated control sequences. First, we reviewed the literature to identify all bacterial strains that have been shown to produce hopanoids in pure culture and thus must encode the SHC enzyme. We also inferred whether HpnP, HpnR, or HpnH also were encoded by these strains based on identification of 2Me-, 3Me- or C_35_ hopanoids, respectively (**Table S1**). Though other hopanoid biosynthesis enzymes that are involved in C_35_ hopanoid modification (HpnG, HpnI, HpnJ, HpnK, and HpnO) have associated profile HMMs, there are few experimental studies on C_35_ sidechain diversity across bacteria. Because we could identify few experimentally validated genes for these enzymes, we excluded them from benchmarking.

We collected genomes for all hopanoid-producing strains in list **Table S1**, as available, and we identified their SHC, HpnP, HpnR, and HpnH homologs by BLAST using hopanoid biosynthesis proteins validated by knockout (KO) or *in vitro* (IV) study as queries. These homologs were assembled into a “positive control” sequence dataset (**Table S2**). KO/IV-verified sequences were also used to query the manually reviewed Swiss-Prot reference collection to generate a “negative control” dataset containing members of related but functionally distinct gene families. We used these control sequences to benchmark the identification of SHC, HpnH, HpnP, and HpnR using each of their associated profile HMMs in **Table 1**. For HpnP, we also evaluated annotation via a combination of the Pfam vitamin B12-binding (PF02310), radical SAM (PF04055), and DUF4070 (PF13282) domains, following the approach used in Hoshino et al.^14^. In the first stage of benchmarking, we assessed whether each profile could successfully identify all and only the positive control sequences (**Table S2**). We also examined each profile’s accuracy by collecting a diverse set of domain-associated sequences in public protein databases, aligning these sequences with our control datasets, and generating maximum likelihood trees (**Figures S1-S4**). These trees allowed us to assess the extent to which proteins annotated with each domain are related to verified sequences versus negative control sequences from families with distinct functions.

All four SHC-associated domains (PF13249+PF13243, TIGR01507, IPR006400, and K06045) were found in >99% of the positive controls, but these domains were also assigned to terpene cyclase negative controls (**Table S2**). In phylogenetic trees of domain-associated proteins, an average of 26.5% of proteins annotated with SHC-related domains are positioned within negative control branches (**Figure S1**). Most of these false positive sequences are homologs of the *Bacillus subtilis* sporulenol synthase, a squalene cyclase that does not produce hopanoids and thus does not belong in the SHC class. With the exception of TIGR01507, the SHC-related domain profiles also cannot distinguish squalene-hopene cyclases from the oxidosqualene cyclases that produce sterols.

Of the hopanoid-modifying enzymes, HpnH-associated domain profiles were most reliable. Both the TIGR03470 and IPR017833 profiles for HpnH detected all positive control sequences and excluded all negative control proteins (**Table S2**). This high accuracy was further supported by phylogenetic trees; proteins annotated with either domain all are within branches including at least one positive control, with no annotated proteins positioned in the negative control branch (**Figure S2A-B**). Domain profiles for the methylases HpnP and HpnR yielded more variable results. Among HpnP-associated domains, the overly stringent K22705 term matched only a minority (32%) of positive controls (**Table S2**); this stringency is also evident in the phylogenetic tree (**Figure S3A**), where most of the positive control sequences are on a separate branch from K22705-annotated proteins. The HpnP annotation method used by Hoshino et al. (PF02310+PF04055+PF13282) failed to capture roughly one third of the positive controls (**Table S2**), and in the phylogenetic tree, proteins annotated with the three Pfam domains do not fully populate the positive control branches (**Figure S3B**). These triply annotated proteins also include HpnP-like proteins that cannot be clearly assigned to either the negative control radical SAM methylases nor true HpnP branches. The IPR034530 domain for HpnP was the only profile we tested that matched all positive and no negative control sequences, though it similarly includes HpnP-like proteins (**Figure S3C**).

Finally, for HpnR, the IPR027564, TIGR04367, and K22704 domains were absent in negative controls, but these profiles did not all detect the positive controls (**Table S2**). We found that the HpnR-associated KO term is the least stringent of the three domain profiles, such that most K22704-annotated proteins do not appear to be true HpnR homologs (**Figure S4A**). Both IPR027564 and TIGR04367 profiles also match numerous uncharacterized proteins in a branch with NirJ proteins (**Figure S4B-C**).

Collectively, our results demonstrate that domain assignments based on pre-computed profile HMMs alone are not a reliable metric for hopanoid gene annotation, and that the source of the optimal domain profile (Intepro/Pfam, TIGRFAM, Kegg Ontology, *etc*.) varies on a case-by-case basis. These findings led us to refine our strategy for identification of hopanoid biosynthesis genes families across bacterial genomes: (i) select HMM profiles that match 100% of the positive control sequences, using the most comprehensive set of experimentally validated sequences possible; (ii) apply these profiles to the genome set; and (iii) remove false positive sequences based on their similarity to positive vs. negative control sequences, as determined using phylogenetic tools.

### Hopanoid biosynthesis genes are distributed across diverse bacterial phyla

To assess hopanoid distribution across bacteria, we collected complete bacterial genomes in the Joint Genome Institute’s Integrated Microbial Genomes (IMG) database^16^, filtered them by quality, and classified them into taxa as described in the Methods. This dataset included 15,541 unique species and a total of 62,143 genomes (**Table S3**). Within each species, we calculated the percentage of genomes encoding SHC, HpnH, HpnP, and HpnR, using the benchmarked annotation approach described in the previous section (**Tables S4-S5**); phylogenetic trees used to identify and remove false positives are shown in **Figure S5**. We also annotated the genomes with C_35_ hopanoid-modifying enzymes HpnG, HpnI, HpnJ, HpnK, and HpnO based on presence of their associated TIGRFAM domains (**Table 1**), though we note that these annotations were not benchmarked for reasons outlined above.

By this metric, we calculated that ∼10% (1645/15541) of species within our dataset have the genetic capacity to produce hopanoids (**Table S4**; **Figure 1**). SHC-containing species are most abundant within the pseudomonadota, with high conservation within multiple families of alphaproteobacteria and the clade containing *Burkholderia* and *Paraburkholderia* spp. (classified as gammaproteobacteria in the Genome Taxonomy Database (GTDB)^17^ and as betaproteobacteria in the List of Prokaryotic names with Standing in Nomenclature (LPSN)^18^). SHC is also highly conserved within acidobacteriota and the GTDB desulfobacterota (a.k.a.deltaproteobacteria and thermodesulfobacteriota in LPSN), as well as within the cyanobacteriota and actinomycetota. Lower levels of SHC conservation are found in planctomycetota, verrucomicrobiota, and chloroflexota. SHC is absent or extremely rare within bacillota, bacteroidota, campylobacteriota, deinococcota, fusobacteriota, synergistota, spirochoaetota, and thermotogota.

**Figure 1.**
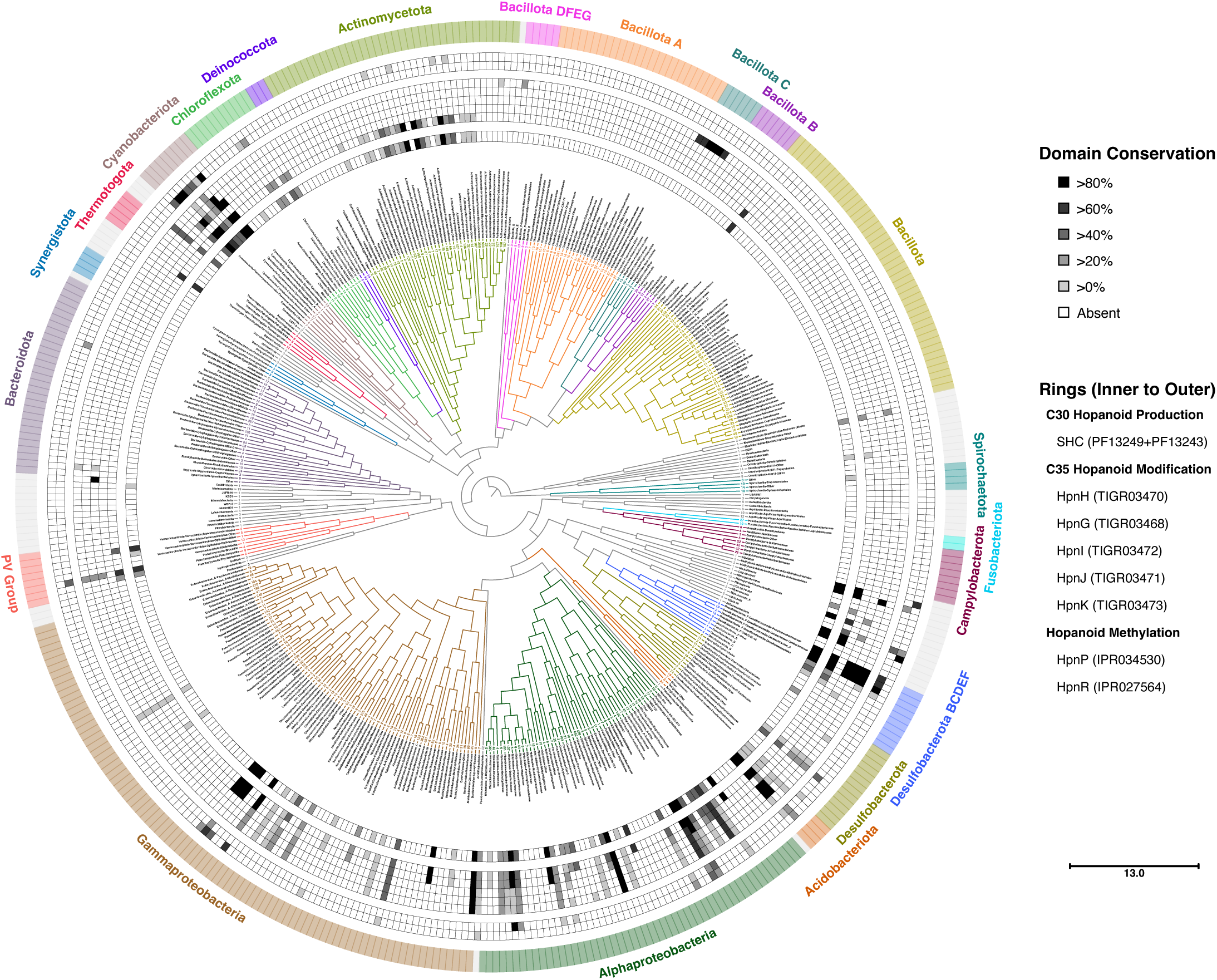
Conservation of hopanoid biosynthesis in bacteria. Maximum likelihood cladogram of bacterial taxa using bac120 sequences. Each branch represents a group of phylogenetically related organisms; the number of genomes represented by each branch and the taxonomic classification of those genomes are provided as branch tip labels and branch names, respectively. Phyla assigned to genomes in each branch using the GTDB-Tk program are shown in the outer ring, with corresponding color-coding of the individual branches. Inner rings with grayscale rectangles indicate the degree of conservation within each branch of various hopanoid-associated domains (see legend for domain/pathway represented by each ring). Scale bar indicates substitutions per 1000 nucleotide positions.

Of the hopanoid-modifying enzymes, HpnH is most common; nearly all SHC-containing genomes also encode HpnH. Domains for HpnG, HpnI, and HpnJ were only identified in genomes that contain the SHC and HpnH domains required for the first steps of C_35_ hopanoid production. HpnK also is involved in downstream modification of C_35_ hopanoids, but its distribution includes some likely spurious annotations within the bacillota. We infer from these distributions that C_35_ hopanoid production and modification is most common in the pseudomonadota, desulfobacterota, acidobacteriota, and cyanobacteriota. Actinomycetota appear to have the capacity to produce ribosylhopane via HpnH but lack the enzymes for downstream modification.

Hopanoid methylases HpnP and HpnR have the sparsest taxon distributions. HpnP and HpnR are both present in pseudomonadota and desulfobacterota, whereas only HpnP is found in cyanobacteriota and planctomycetota, and only HpnR is present in actinomycetota. We note that within cyanobacteria, conservation of HpnP is not universal. Within the cyanobacteriia class – which contains all of the complete, high quality cyanobacterial genomes in IMG – hopanoid production is absent in both the synechococcales order (dominated by *Synechococcus elongatus* genomes) and the PCC-6307 order (dominated by *Prochlorococcus* spp. genomes) (**Table S5**).

These results differ from the HpnP distribution in cyanobacteria reported by Hoshino *et al*.^14^. We hypothesized this discrepancy could arise from differences in the genomes examined, the taxonomic classification of those genomes, or in our HpnP identification strategies. We assessed these hypotheses by collecting all cyanobacterial genomes evaluated in the other study, assigning taxa to those genomes using GTDB-Tk, and re-calculating their HpnP conservation levels using the other group’s HpnP identification strategy. We also evaluated their list of cyanobacterial HpnP proteins against our benchmarking strategy. When using the same taxon classification strategy, our results generally agree (**Table S6**); our observation that HpnP homologs are absent in synechococcales arises from the re-classification of many genomes annotated as synechococcales members in LPSN as members of the thermosynechococcales order in GTDB. When HpnP is present, we do observe higher levels of conservation than the other study, which we attribute to the limitations of the other group’s HpnP identification strategy (co-annotation with PF02310+PF04055+PF13282). This is supported by our phylogenetic analyses of the HpnP sequences identified by Hoshino *et al.*^14^, which do not capture the full diversity of verified HpnP sequences, even when comparing only to the subset of experimentally verified HpnP homologs from cyanobacteria (**Figure S6**).

### Hopanoid production likely arose in marine alphaproteobacteria

Our genome analysis (**Figure 1**) suggested two possible origins for hopanoid biosynthesis: an ancestor of the clade containing the cyanobacteriota, chloroflexota, deinococcota, and actinomycetota, or an acestor of the clade containing the pseudomonadota, acidobacteriota, and desulfobacterota. To assess these possibilities, we generated maximum likelihood phylogenetic trees containing all SHC homologs in our genome dataset identified the most likely root of the tree by rootstrapping. We also assessed the origins of C_35_ hopanoid production, 2Me-hopanoid production, and 3Me-hopanoid production by generating trees for HpnH, HpnP, and HpnR.

The maximum likelihood phylogenetic tree for SHC homologs (**Figure 2**) suggests that SHC originated in marine alphaproteobacteria, including members of the parvularculaceae and CGMCC-115125 (*Pyruvatibacter* spp.) families, and was present in other (non-marine) aquatic alphaproteobacterial families before arising in other lineages. From these aquatic alphaproteobacteria, SHC proteins form three distinct clades. One clade includes terrestrial alphaproteobacteria and a subset of SHC-encoding burkholderiaceae, whereas the SHC proteins from the majority of hopanoid-producing gamma-proteobacteria form a separate clade originating in the marine nitrococcaceae and salinisphaeraceae families. All non-proteobacterial SHC proteins are present in a separate clade that originates in the newly cultivated methylaequoraceae/TMED127 family of alphaproteobacteria, including the SHC proteins from desulfobacterota, acidobacteriota, cyanobacteria, and actinobacteria.

**Figure 2.**
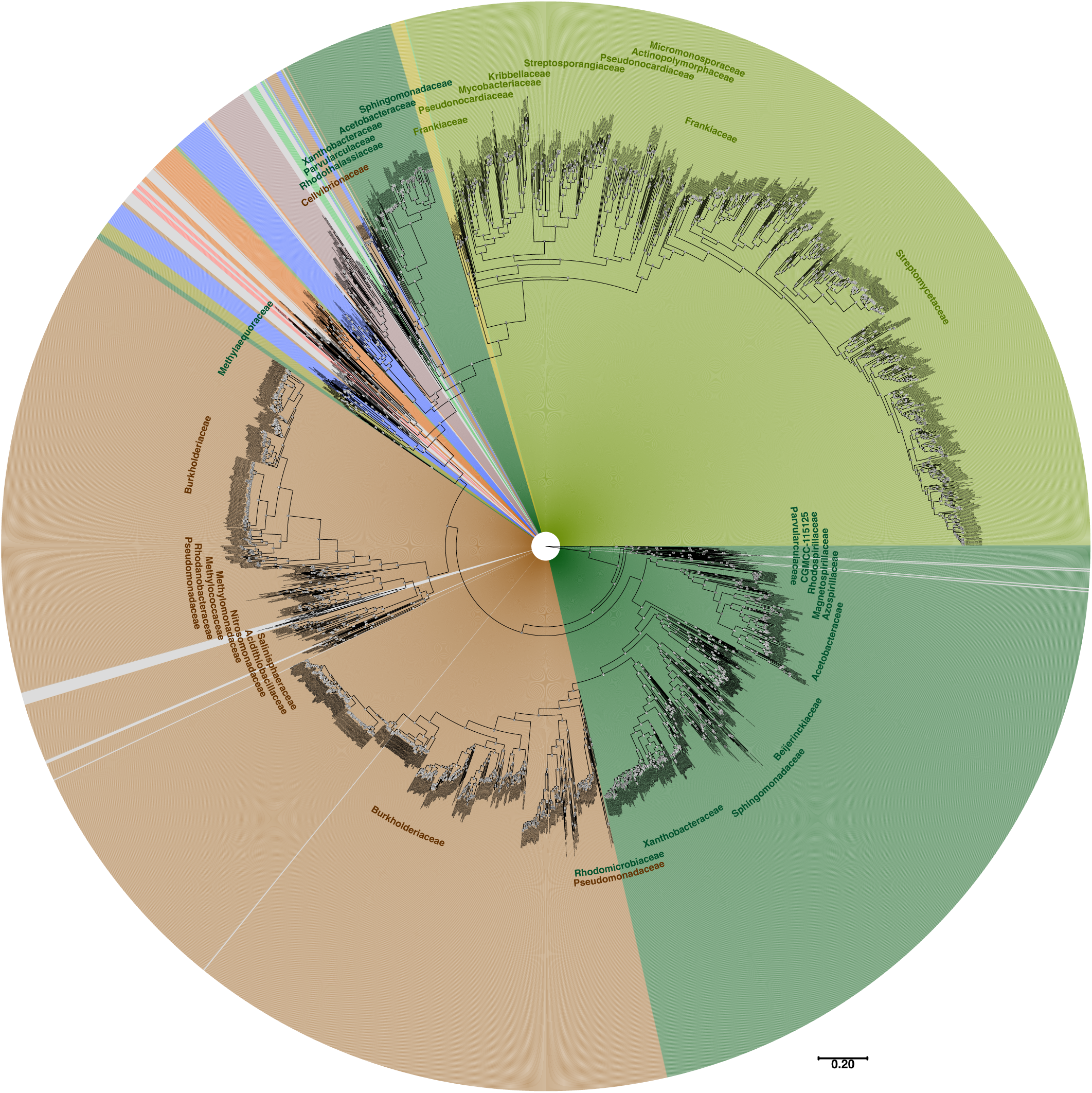
Hopanoid biosynthesis originated in marine alphaproteobacteria. Maximum likelihood phylogenetic tree generated for protein sequences associated with SHC. Branch names include IMG genome ID, GTDB family and genus assignments, and IMG protein ID. Numbers at internal nodes (highlighted in white) indicate branch support values using ultrafast bootstrap approximation (UFBoot), based on 1000 bootstraps; 100 = highest confidence, 0 = no confidence. Scale bar indicates substitutions per 1000 amino acids. Branch backgrounds are color-coded by phyla according to the color scheme (outer ring and phylum text labels) used in Figure 1.

The maximum likelihood tree of HpnH homologs is surprisingly similar (**Figure S7**), containing clades with similar composition to the SHC tree that also appear to originate in the parvularculaceae, CGMCC-115125, and methylaequoraceae families of marine alphaproteobacteria. Given the similarity of the SHC and HpnH trees and their highly overlapping distributions, we conclude that the capacity to produce C_35_ hopanoids occurred early in the evolution of hopanoid biosynthesis. That the roots of both SHC and HpnH trees were identified the same families of organisms provides strong evidence – beyond the individual trees’ root confidence values (>85% in both cases) – that hopanoids originated in marine alphaproteobacteria.

The origins of methylated hopanoids are less clear. In the maximum likelihood tree for HpnP proteins, two possible origins are suggested: one major clade potentially originating in the acidobacteriota (though we note our dataset contained few acidobacterial proteins), from which the cyanobacterial and proteobacterial HpnP proteins evolved, and a minor clade originating in the desulfobacterota containing HpnP proteins from all other 2Me-hopanoid-producing phyla (**Figure 3**). However, this root location is weakly supported by rootstrapping. Statistical analyses of all possible root positions via tree topology tests identified 37 potential root positions resulting in trees with a statistically similar likelihood (p > 0.05 threshold) as the maximum likelihood tree (**Dataset S1**). Interestingly, none of these trees suggest that 2-Me hopanoids originated in cyanobacteria. A plurality of likely root positions indicate that HpnP first appeared in both the acidobacteriota and the desulfobacterota, and alternative origins in the planctomycetota or chloroflexota were also common. However, we cannot rule out long-branch effects as the explanation for these potential root positions, especially given the paucity if *hpnP* sequenced from these phyla.

**Figure 3.**
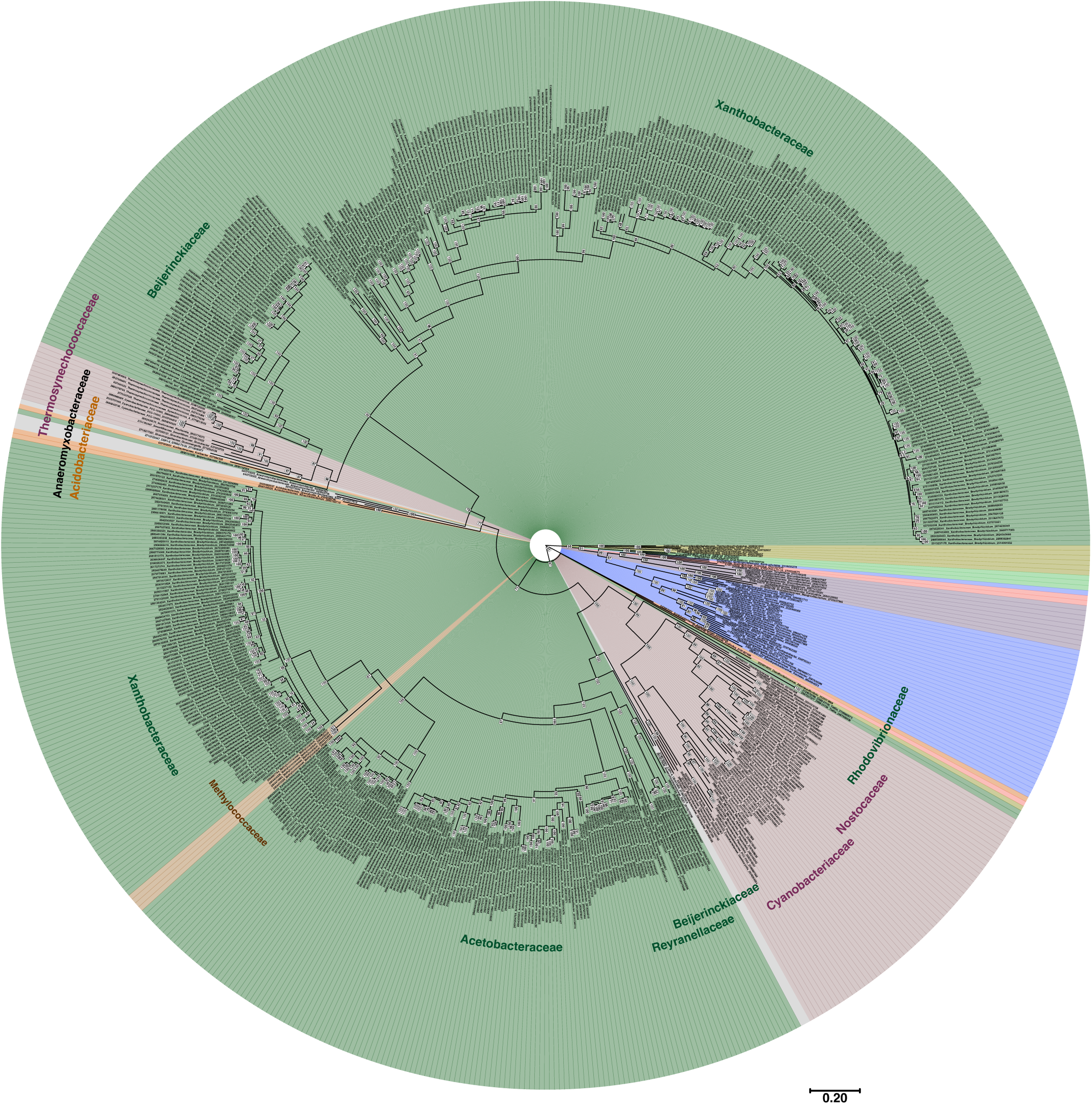
Phylogenetic relationships of HpnP homologs. Maximum likelihood phylogenetic tree generated for protein sequences associated with HpnP. Branch names include IMG genome ID, GTDB family and genus assignments, and IMG protein ID. Numbers at internal nodes (highlighted in white) indicate branch support values using ultrafast bootstrap approximation (UFBoot), based on 1000 bootstraps; 100 = highest confidence, 0 = no confidence. Scale bar indicates substitutions per 1000 amino acids. Branch backgrounds are color-coded by phyla according to the color scheme (outer ring and phylum text labels) used in Figure 1.

The origin of 3Me-hopanoids is also ambiguous. The maximum likelihood tree of HpnR sequences tree indicates that HpnR proteins may have evolved from a shared ancestor of pseudomonadota and acidobacteriota (**Figure S8**) and were transferred to actinomycetota through these groups. The HpnR proteins from zetaproteobacteria, myxococcota, and methylomirabilota may have originated independently as they are present on a separate clade from the majority of HpnR sequences. Because the predicted root position of the tree is weakly supported by rootstrapping, we also performed tree topology tests as described for HpnP (**Dataset S1**). The 17 equally likely root positions (p > 0.05 threshold) generally agree with interpretations based on the maximum likelihood tree, though some root positions nest the zetaproteobacteria, myxococcota, and methylomirabilota HpnR proteins within one of the two major clades.

### Hopanoid biosynthesis genes are correlated with nitrogen cycling and high osmolarity tolerance

To find shared features of hopanoid-producing bacteria, we identified all protein domain families that are enriched within genomes encoding SHC and other hopanoid biosynthesis genes (**Table S7**). This analysis revealed several protein functional categories that are positively correlated with hopanoid production (**Table 2**). The most enriched domains in SHC-containing genomes were associated with synthesis and modification of hopanoids, as well as of squalene, isoprenoid precursors (farnesyl and geranyl diphosphate), and other cyclic isoprenoids such as carotenoids or sterols. An enrichment of domains associated with production of B vitamins and S-adenosyl-methionine (SAM) metabolism also was apparent, likely relating to the role of vitamin B12 and SAM as co-factors for hopanoid biosynthesis proteins HpnH, HpnP, and HpnR.

**Table 2.** Protein domain categories positively correlated with SHC. Manually annotated domain categories correlated with SHC across all genomes. Center column represents average correlation coefficient (r) for domains within each category. A full list of correlated domains is provided in **Table S7**.

| Terpenoid/isoprenoid biosynthesis & co-factors |  |  |
| --- | --- | --- |
| *Function* | *r* | *Examples* |
| **Hopanoid synthesis** | 0.595 | squalene-hopene cyclase, hopanoid biosynthesis associated radical SAM protein HpnH, hopanoid C-2 methylase, hopanoid C-3 methylase |
| **Squalene synthesis** | 0.412 | presqualene diphosphate synthase, squalene synthases HpnC & HpnD, hydroxysqualene synthase |
| **Radical SAM enzymes** | 0.225 | methylthioribose-1-phosphate isomerase, S-adenosyl-L-homocysteine hydrolase, ergothioneine biosynthesis proteins EgtABC, coenzyme PQQ biosynthesis enzyme PqqE |
| **Other terpenoid/isoprenoid synthesis** | 0.217 | farnesyl-diphosphate farnesyltransferase, geranyl diphosphate 2-C-methyltransferase, 2-methylisoborneol synthase, germacradienol/geosmin synthase, xanthine dehydrogenase, menaquinone-9 beta-reductase |
| **B vitamin synthesis** | 0.195 | precorrin-3B synthase, cobalamin biosynthesis protein CobT, thiamine biosynthesis protein ThiS, phosphoribosylformylglycinamide (FGAM) synthase |
| **Sterol metabolism/transport** | 0.176 | cholesterol oxidase, cholesterol uptake porter CUP1 of Mce4, phospholipid/cholesterol/gamma-HCH transport system proteins |
| Cellular homeostasis pathways |  |  |
| *Function* | *r* | *Examples* |
| **Trehalose/starch synthesis** | 0.248 | malto-oligosyltrehalose synthase, trehalose synthase, starch synthase (maltosyl-transferring), maltose alpha-D-glucosyltransferase / alpha-amylase |
| **Osmoresponsive signaling** | 0.203 | osmosensitive K<sup>+</sup> channel His kinase sensor domain, K<sup>+</sup>-transporting ATPase (KDP operon) subunits, KDP operon response regulator KdpE and sensor histidine kinase KdpD |
| **Divalent cation transport** | 0.198 | copper resistance proteins, periplasmic divalent cation tolerance protein, HoxN/HupN/NixA family nickel transporter, high affinity Mn<sup>2+</sup> porin |
| **Saccharide transport/modification** | 0.194 | beta-1,3-glucanase, beta-glucan exporter, alpha-glucan phosphorylases, non-reducing end alpha-L-arabinofuranosidase, isoamylase, OprB family carbohydrate-selective porin, xylobiose transporters |
| Taxon-specific metabolisms |  |  |
| *Function* | *r* | *Examples* |
| **Single C, N source utilization** | 0.213 | ammonia/methane monooxygenase, carbon-monoxide dehydrogenase |
| **Pentose phosphate pathway** | 0.205 | 6-phosphogluconate dehydrogenase, xylulose-5-phosphate/fructose-6-phosphate phosphoketolase, fructose-1,6-bisphosphatase II / sedoheptulose-1,7-bisphosphatase |
| **Methylotrophy** | 0.196 | formylmethanofuran dehydrogenase, methenyltetrahydromethanopterin cyclohydrolase, Cof coenzyme F420 synthases |
| **Aerobic respiration** | 0.195 | NADH:quinone oxidoreductase, cytochrome cbb3 oxidase, Cta/Caa/Cox-type cytochrome c oxidases |
| **Glyoxylate pathway** | 0.195 | glycolate oxidase, phosphoenolpyruvate guanylyltransferase, phosphoenolpyruvate phosphomutase |
| **Nitrogen fixation** | 0.177 | nif-specific ferredoxin, nitrogen fixation proteins NifN, NifT, NifW, NifX, FixT, FixU |

The second most significant category of correlated proteins includes those involved in osmotic pressure sensing and response. Domains involved in biosynthesis of the osmolyte trehalose and other saccharides involved in osmotic stress (*e.g.*, osmoregulated periplasmic glucans) are enriched in hopanoid-producers to a similar level as other isoprenoid biosynthesis domains. This category also includes the osmotic stress-regulated two-component system KdpPE, which controls potassium flux across the membrane via the potassium pump KdpFABC, and various transporters involved in divalent cation homeostasis. Interestingly, osmo-responsive protein domains are correlated with SHC across nearly all hopanoid-producing taxa.

Specific bacterial metabolisms were also highlighted by our analysis. Domains involved in utilization of single-carbon substrates and in nitrogen cycling are common to SHC-encoding genomes, and the biggest group of SHC-correlated metabolic domains are involved in aerobic respiration, such as the high efficiency terminal oxidases required for nitrogen fixation. To test this relationship between SHC and the nitrogen cycle, we measured the growth of a hopanoid-deficient strain of *Bradyrhizobium diazoefficiens*^19,20^ on several nitrogen cycle intermediates (**Figure S9**). We find that hopanoid-deficient mutants grow slowly in the presence of nitrite but have no growth defect in the presence of ammonia or nitrate. Our results support other recent work linking hopanoids to nitrite metabolism^10–12^.

### Microsynteny suggests novel hopanoid transport factors

Roughly 16% of the top 1000 domains positively correlated with SHC were annotated as uncharacterized or domains of unknown function (DUFs) (**Table S7**); collectively, this group was the most enriched category in SHC-containing genomes. We hypothesized that any DUFs directly involved in hopanoid biosynthesis would be adjacent to other hopanoid-associated genes within the genome. To identify such DUFs, we performed a targeted analysis of protein domains found near the *shc* gene locus. We collected domain annotations for all genes within 20 ORFs up- or down-stream of *shc* in each genome, categorizing these genes into families based on shared domain annotations (**Table S8**).

Our analysis revealed four uncharacterized gene families that are commonly located near the *shc* locus and have potentially hopanoid-related functions (**Figure 4**). Two of these families are annotated with domains that already were computationally predicted to be related to hopanoids. The *hpnL* gene family includes genes annotated with the uncharacterized “HpnL” domain in TIGRFAM (TIGR03476); this domain does not exist in other protein family or pathway databases, and in InterPro is instead annotated as a member of the lysylphosphatidylglycerol synthetase/glycosyltransferase AglD (IPR022791) family. The AglD InterPro domain includes proteins involved in addition of lysine addition to the phospholipid phosphatidylglycerol and in microbial S-layer modification. The *hpnM* gene family is annotated with the InterPro “Hopanoid biosynthesis-associated membrane protein HpnM” domain (IPR017842), a subset of “Toluene tolerance Ttg2/phospholipid-binding protein MlaC” (IPR008869) domain proteins. Two new candidate hopanoid gene families, which we name here *hpnS* and *hpnT*, have not previously been associated with hopanoids. The *hpnS* family genes share the domain annotation MlaA lipoprotein (IPR007428). The *hpnT* genes are annotated with Domain of unknown function DUF2147 (IPR019223); a crystal structure of the DUF2147 domain-containing protein HP1028 from *Helicobacter pylori*^21^ suggests these domains belong to the lipocalin family, a group of proteins that transport hydrophobic small molecules^22^.

**Figure 4.**
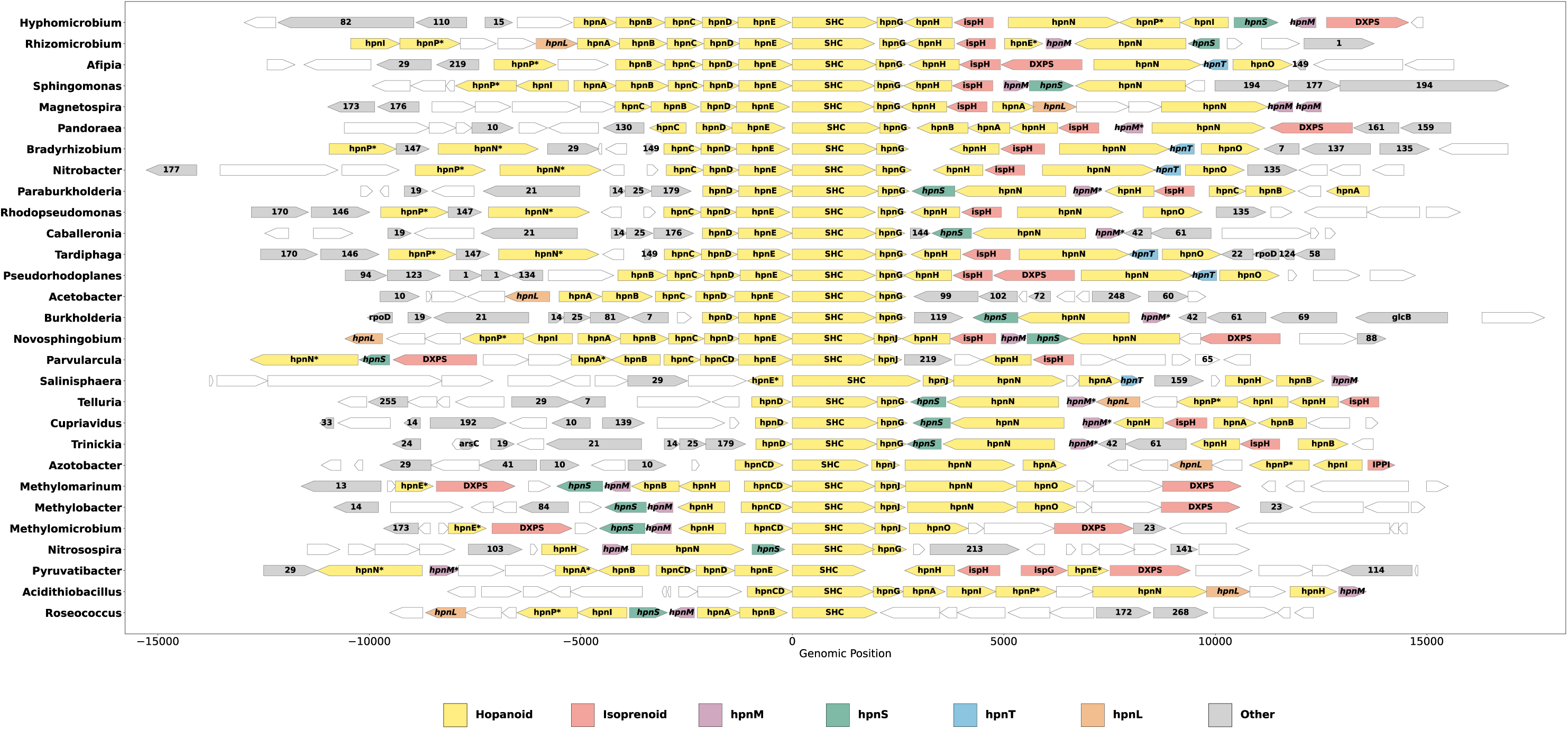
Candidate hopanoid-associated genes *hpnL*, *hpnM*, *hpnS*, and *hpnT* are enriched near *shc* loci. Genes adjacent to the *shc* locus in selected bacterial genomes. Color-coding of ORFs indicates functional categories for conserved gene families: known hopanoid-associated genes (Hopanoid, yellow); known isoprenoid biosynthesis genes (Isoprenoid, pink), and candidate gene families *hpnM*, *hpnS*, *hpnT*, and *hpnL* (purple, green, blue, and orange, respectively). ORFs in the Other category (grey) are conserved gene families with unknown functions; number labels correspond to specific protein domain annotations provided in Table S3. Gene names labeled with asterisks indicate likely gene famly members that lack one or more domain annotation relative to other genes in the same family (*e.g.,* ‘*hpnM*\**’* = *‘hpnM-like*’). All unlabeled ORFs (white) are non-conserved genes.

In *E. coli* and other diderms, MlaC and MlaA are components of an intermembrane phospholipid transport system^23,24^, whereas lipocalins can serve as extracellular lipid carriers in eukaryotes^25^. We thus hypothesized that HpnM, HpnS, and HpnT may form components of inner-to-outer membrane transport systems for hopanoids, in concert with the inner membrane hopanoid transporter HpnN. This possibility is supported by the restriction of these proteins to diderms and the presence of predicted signal peptides, indicating a likely periplasmic localization (**Figure S10**). Our subsequent analysis of the conservation of *hpnM*, *hpnS*, and *hpnT* genes within the vicinity of *hpnN* loci (**Table S8**) suggests that they evolved in distinct taxa: the lipocalin-like HpnT protein is conserved in the hyphomicrobiales/rhizobiales order of alphaproteobacteria, whereas both HpnM and HpnS co-occur in the sphingomonadales, rhodospirillales, and azospirillales orders of alphaproteobacteria and across gammaproteobacteria (**Figure 5**).

**Figure 5.**
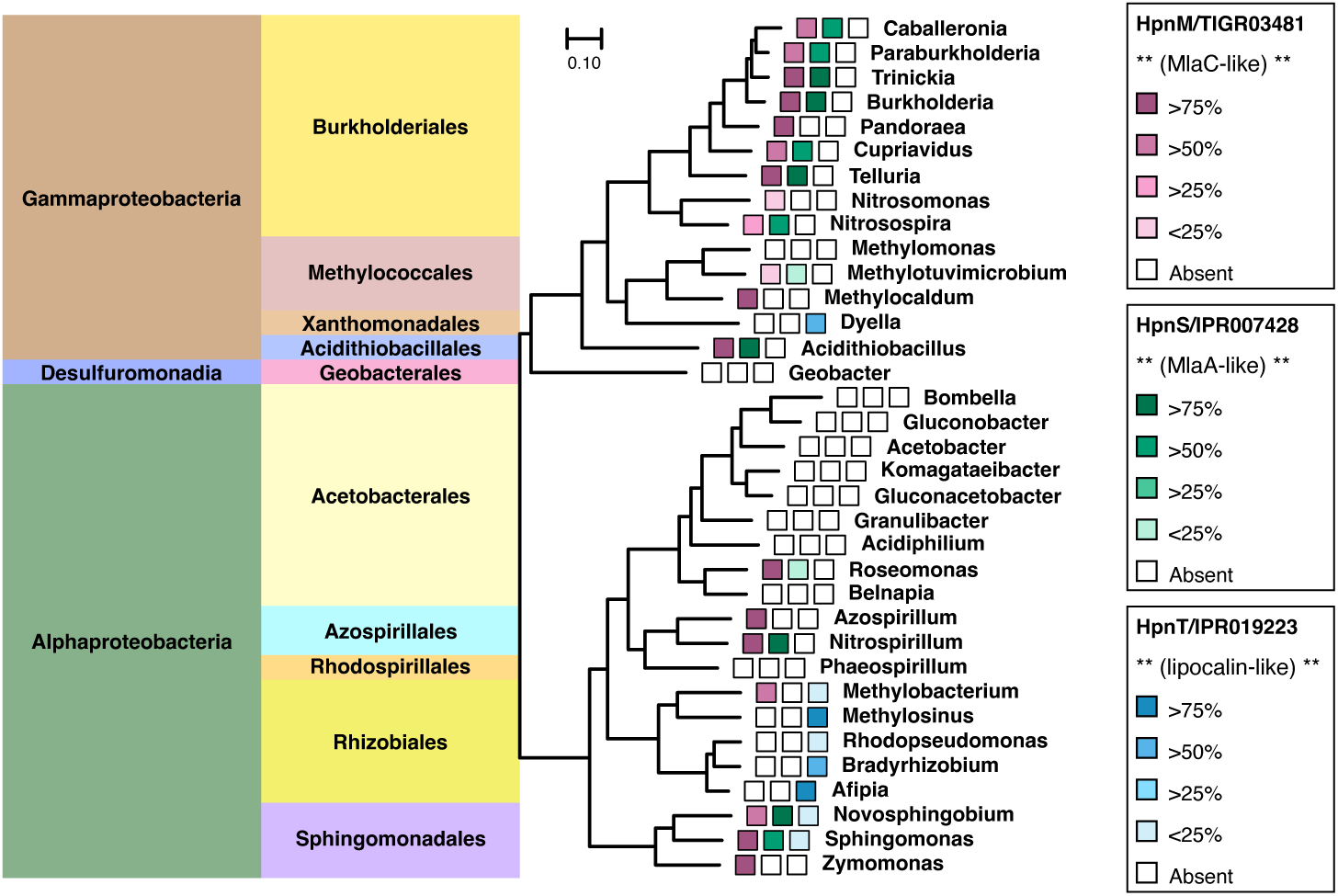
The HpnM/S module and HpnT appear in distinct HpnN-encoding taxa. Maximum likelihood tree of bac120 sequences from selected HpnN-encoding, Gram-negative bacterial genera. Branch tip labels (shaded rectangles) indicate conservation of HpnM, HpnS, and HpnT across all HpnN-encoding genomes within each genus; conservation levels corresponding to each shade are provided in the legend. Presence of these proteins was not annotated in the archaeon outgroup, *Thermococcus*. Scale bar indicates substitutions per 1000 nucleotides.

MlaFEDB proteins form a complex^26^ that moves phospholipids through the inner membrane, delivered by the periplasmic shuttle protein MlaC, which in turn ferries phospholipids from the MlaA lipoprotein that associates with outer membrane proteins^24,27,28^ (**Figure 6A**). We speculated that HpnM and HpnS might also form a shuttle and OM anchor protein system, which would require them to bind one another at the OM (**Figure 6B**). To test this hypothesis, we used AlphaFold 3^29^, which predicted a high probability that the *Burkholderia multivorans* HpnM protein binds both the hopanoid diploptene and HpnS in a manner similar to the predicted MlaA/MlaC dimer^30^ (**Figure 6C**). *Bradyrhizobium diazoefficiens* HpnT is predicted to bind diploptene and may also act as a periplasmic shuttle, though it has no obvious cognate OM binding partner. These structural predictions support the hypothesis that HpnM, HpnS, and HpnT are novel hopanoid trafficking factors.

**Figure 6.**
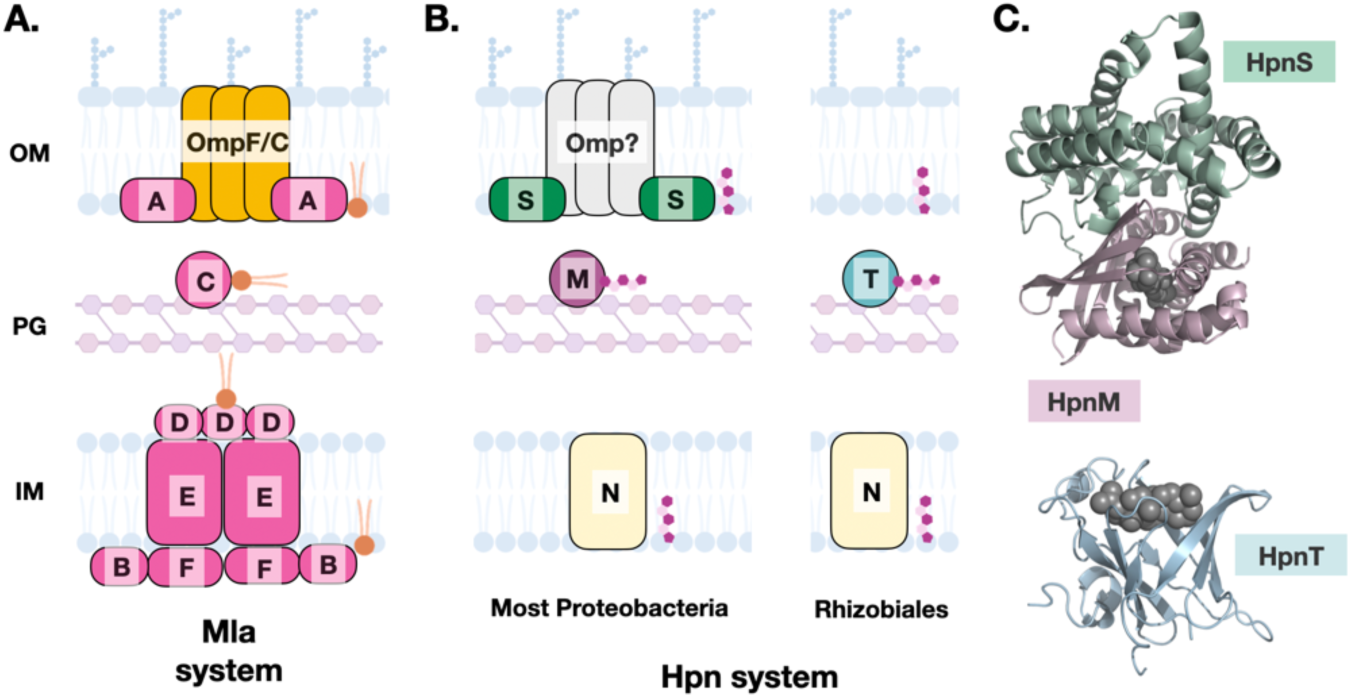
Candidate hopanoid transport pathways in proteobacteria. (**A**) Schematic of the Mla phospholipid (PL) transport system from *E. coli*. The inner membrane transport complex MlaBDEF transports PLs to the periplasmic shuttle protein MlaC, which interacts with OmpF/C-associated lipoprotein MlaA in the outer membrane. (**B**) Candidate hopanoid transport pathways in diderm bacteria. In most proteobacteria (left), the inner membrane transporter HpnN may interact with the periplasmic shuttle HpnM to deliver hopanoids to the outer membrane lipoprotein HpnS. In the Rhizobiales order (right), hopanoids trafficked by HpnN are delivered to the periplasmic HpnT protein. (**C**) AlphaFold 3-predicted structures of the *Burkholderia multivorans* HpnSM dimer (top) and *Bradyrhizobium diazoefficiens* HpnT (bottom) in association with the hopanoid diploptene (grey).

## MATERIALS & METHODS

### Assembly of control sequence datasets

Strains experimentally verified to produce hopanoids were identified by literature review, and strain names were updated manually to their preferred LPSN names by searching the strains’ culture collection IDs in the DMZ StrainInfo database^31^. The LPSN strain names were used to identify corresponding genome assemblies from NCBI. Homologs of SHC, HpnH, HpnP, and HpnR were identified in these genomes by BLAST search (e-value < 1e-50) using knockout- or biochemically-verified enzymes as queries. Genome accession numbers and relevant citations for these queries are indicated with asterisks in **Table S1**, and the accession numbers of corresponding BLAST hits are provided in **Table S2**. Negative control sequences were determined by BLAST search of the manually curated Swiss-Prot database using knockout- or biochemically-verified SHC, HpnH, HpnP, and HpnR queries. Selected negative control sequences (accessions provided in **Table S2**) represent the top BLAST hits with experimental evidence to demonstrate that they are involved in functions distinct from hopanoid biosynthesis.

### Protein domain profile benchmarking

Protein domain profiles matching negative and positive control sequences were determined using the InterProScan command line tool (version 5.69-101.0, https://github.com/ebi-pf-team/interproscan)^32^ for Pfam, InterPro, and TIGRFAM domains and KofamKOALA (ver. 2026-01-01, KEGG release 117.0, https://www.genome.jp/tools/kofamkoala/) ^33^ for Kegg Ontology domains. For phylogenetic analysis-based benchmarking, all proteins matching each SHC, HpnH, HpnP, and HpnR-associated domains in **Table 1** were collected from UniProt (for Pfam and InterPro domains) or Joint Genome Institute’s Integrated Microbial Genomes (IMG) database (https://img.jgi.doe.gov/)^16^(for TIGRFAM and Kegg Ontology domains). Matching proteins were filtered by similarity using a 75% identity cutoff CD-Hit (cd-hit, https://github.com/weizhongli/cdhit/wiki)^34^ and were aligned, along with their corresponding positive and negative control sequences, using the *super5* algorithm of the MUSCLE v. 5.1 sequence alignment program (https://www.drive5.com/muscle/muscle.html)^35^. This alignment was used to generate a maximum likelihood phylogenetic tree with IQ-TREE 2 (https://iqtree.github.io/)^36^ using the Q.pfam amino acid substitution model. Trees were visualized and annotated with control sequence positions using TreeViewer^37^.

### Bacterial genome selection

Genomes were sourced from IMG and filtered by quality using CheckM2 v1.0.2^38^. We selected only genomes meeting the following criteria: (i) completeness estimate >50%, (ii) contamination estimate <10%, (iii) quality score (completeness - 5*contamination) >50, (iv) <1000 contigs, (v) N50 > 5 kb, and (vi) presence of >40% of the bac120 gene regions used for taxon assignment by GTDB. Genomes passing these criteria were further filtered to remove accessions where <75% of genes were associated with one or more TIGRFAM, Pfam, or Kegg Ontology annotation(s), indicating incomplete annotation. We also removed genomes with a total gene count < 1000, as these genomes are predominantly from obligate endosymbionts with heavily reduced genomes. Complete sequences for accessions that met these criteria (totaling 62,143 genomes, listed in **Table S3**) were downloaded from IMG along with their associated ORF, protein domain, and functional pathway annotations.

### Genome classification

Taxonomic classification of the genomes was performing using the GTDB Toolkit v2.4.0+ (https://github.com/Ecogenomics/GTDBTk)^39^. For the majority of genomes, GTDB-Tk was sufficient to provide annotations through the species level. However, a subset of the genomes did not match one of the GTDB reference strains. For these accessions, tentative species were assigned by 16S rRNA gene similarity, using the canonical 97% sequence identity cutoff using CD-Hit. Because genomes in our dataset contained an average of three 16S genes, 16S-based assignments were not always consistent. In 1.2% of cases, 16S sequences from the same genome did not share at least 97% identity, and in ∼9% of genomes, 16S-based genus assignments did not match assignments based on whole-genome annotations (**Table S9**). We addressed this by discarding taxon assignments from 16S sequences for which the associated genome had a different genus assignment than the associated 16S cluster.

### Generation of bac120 phylogenetic tree

For each species in our dataset, a single genome was selected as a species representative based on completeness of its corresponding bac120 region. These representative bac120 sequences were pre-aligned by GTDB-Tk, and a maximum likelihood phylogenetic tree was generated using IQ-TREE 2. First, the optimal substitution model was determined by ModelFinder^40^, and a maximum likelihood tree using that model was generated using 1,000 replicates each for calculation of branch support values using ultrafast bootstrap approximation (UFBoot)^41^.

To improve readability, species representatives were further consolidated into higher level groupings such that each group contained at least 5 genomes (which the exception of phyla for which fewer than 5 sequenced genomes are available, which remained single groups), resulting in 445 higher order taxon groups. Representatives for the 445 groups were selected based on the number of genomes, and only these representatives were shown on the final bac120 tree in **Figure 1**. This final tree was annotated to include the corresponding taxon for each branch and the number of genomes represented by each branch in TreeViewer.

### Calculation and visualization of hopanoid biosynthesis gene conservation

We annotated the protein-coding sequences of all genomes in our dataset using command-line InterProScan. We then extracted the sequences for all proteins matching the broadest protein domain profiles for SHC (Pfam13249+Pfam13243), HpnH (TIGR03470), HpnP (IPR034530), and HpnR (TIGR04367). These proteins were aligned with the appropriate set of positive and negative control sequences, and a maximum likelihood tree was generated as described in the *Protein domain profile benchmarking* section.

We calculated the percentage of genomes within each species with at least one protein sequence within the positive control branches of the SHC, HpnH, HpnP, and HpnR trees (**Table S4**), respectively, as well as the percentages of genomes annotated with domains associated with the extended hopanoid-modifying proteins listed in **Table 1**. These percentages were recalculated at the order rank (**Table S5**) and the 445 higher order taxon groups described in the previous section by taking the average percentages for all species within each order/taxon group. Percentages were mapped onto the bac120 tree and visualized using TreeViewer.

### Protein tree generation

For protein trees, the SHC, HpnH, HpnP, and HpnR sequences identified above were aligned without control or outgroup sequences in MUSCLE. For each protein family, we generated 500 diversified alignments using the *align* algorithm, each with a unique seed, and the best scoring alignment was identified by maximum column confidence score using *maxcc*. To increase downstream processing time, columns in which >95% of sequences contained gaps were removed from the best-scoring alignment prior to tree generation.

Maximum likelihood trees were generated using IQ-TREE 2 with 1,000 bootstrap replicates using UFBoot and root positions were inferred using the general non-reversible amino acid model NON-REV^42^. For maximum likelihood trees with low rootstrapping support values (<75%), tree topology tests were performed using tree topology tests. Possible root positions were identified as roots from trees with statistically similar (p > 0.05) likelihoods as the maximum likelihood tree using at least one of the following significance tests: one sided Kishino-Hasegawa test, Shimodaira-Hasegawa test, Expected Likelihood Weight (Strimmer & Rambaut), approximately unbiased (AU) test^42^. As above, all maximum likelihood trees were visualized and annotated using TreeViewer.

### Species tree – gene tree reconciliation for HpnP

A distribution of 1,000 gene trees for HpnP was generated using MrBayes v3.2.7a^43^ using a mixed amino acid substitution model for 1 million generations; all other parameters were set to default values. A corresponding species tree was generated in IQ-TREE 2 using aligned bac120 sequences from GTDB-Tk. The tree included bac120 sequences from all HpnP-encoding species as well as group representatives from Figure 1, which served as outgroups. The root of the species tree was inferred by rootstrapping as described in the previous section. Reconciliation was performed with AleRax v1.4.0 using all 1,000 gene trees for calculation of duplication, transfer, and loss (DTL) probabilities. Predicted DTL events with >50% probability were extracted from AleRax and used to annotate the HpnP species tree in TreeViewer via custom Phyton scripts.

### Calculation of protein domain correlations

We calculated correlation coefficients between SHC, HpnH, HpnP, HpnR and all other bacterial protein domains by first determining whether each Pfam, TIGRFAM, KO, and InterPro domain was present in the genome of each species representative. We then calculated the Pearson correlation coefficient and corresponding p-value for every pair of domains using binary presence (1) and absence (0) values, either across all genomes in the data or within phyla (**Table S7**). Correlation coefficients were calculated in custom Python scripts using the *scipy.stats.pearsonr* function.

### B. diazoefficiens nitrogen utilization assays

Wild-type Bradyrhizobium diazoefficiens strain USDA110 spc4 and inducible hopanoid mutant strain Pcu-shc::Δshc (abbreviated as Δshc) were cultured as described previously ^20^. Briefly, an individual colony of each strain was incubated in liquid AG liquid media aerobically in an incubating shaker at 30°C and 250 rpm until mid-log phase (OD_600_ = 0.4–0.8). WT, Δ*shc,* and Δ*shc* with cumate supplementation (Δ*shc* + c) were diluted into AG minimal media lacking a nitrogen source (2.3 mM sodium gluconate, 3.3 mM arabinose, 5.6 mM MES monohydrate, 5 mM HEPES, 1 mM Na2HPO4, 1.76 mM Na2SO4, 88 µM CaCl2, 25 µM FeCl3, and 0.73 mM MgSO4, pH 6.6) to an initial OD_600_ = 0.008. Aliquots of this dilute subculture (100 µl) were dispensed into a 96-well Biolog PM3 Microplates containing 95 different nitrogen sources and a negative control. Each well was overlaid gently with 30 µL of mineral oil to prevent evaporation. The plate was placed in a BioTek microplate reader with continuous shaking, and the OD_600_ was measured every 2 h for 168 h. The growth curve data were plotted using Prism GraphPad.

### Microsynteny analysis of shc- and hpnN-adjacent genomic regions

To compare regions surrounding *shc* across genomes, we selected the subset of genomes we found to contain SHC. We defined *shc*-adjacent regions (“neighborhoods”) as the nearest 20 upstream and 20 downstream neighboring ORFs. We used InterProScan to identify the protein domains encoded by each neighboring gene, and then we assigned each gene a numerical ID based on the set of InterPro domains that it contained. For some genes, such as those known to be involved in hopanoid or isoprenoid biosynthesis, we replaced the numerical ID with a gene name such as “hpnN” or “hpnM”. In some cases, genes contained some but not all of the domains of a known gene, and we assigned these genes the name of the known gene followed by an asterisk (*e.g.*, “hpnN*” or “hpnM*”). InterPro domains that occurred in fewer than 5% of the neighboring genes were excluded from this analysis, and genes containing only those domains were not assigned a name. To visualize microsynteny in these neighborhoods, we obtained the coordinates and orientation of each *shc*-neighboring gene and used a custom Python script to draw the genomic regions. To align the regions, the starting coordinate of *shc* was set to 0 for each genome. We colored each gene based on its assigned name/numerical ID and illustrated gene orientation using triangles pointed toward the 3’ end.

To evaluate genes in the *hpnN* neighborhood, we selected genomes containing *hpnN* and removed any genomes that were not assigned to a Gram-negative species. We then calculated the percentage of genomes with *hpnM*, *hpnS*, and *hpnT* in the *hpnN* neighborhoods across genera. These values were visualized on a maximum likelihood tree of bac120 sequences for species representatives of *hpnN*-containing, Gram-negative genera generated as described in the *Generation of bac120 phylogenetic tree* section and visualized in TreeViewer.

### HpnT, HpnM and HpnS structure and binding predictions

To predict the three-dimensional structures of HpnT, HpnM, and HpnS with diploptene, we used AlphaFold 3 (AF3, https://github.com/google-deepmind/alphafold3)^29^. For these predictions, we used the protein sequences of HpnM and HpnS from *Burkholderia multivorans* ATCC 17616 (BMULJ_05342 / Uniprot A0A0H3KU80 and BMULJ_05340 / Uniprot ID A0A0H3KPM6) and the protein sequence of HpnT from *Bradyrhizobium diazoefficiens* USDA110 (*bll3009*, Uniprot ID Q89QW5). The SMILES string of diploptene was obtained from PubChem. For HpnT, we predicted the structure of a HpnT monomer in complex with diploptene. For HpnM and HpnS, we predicted the structure of a HpnM / HpnS heterodimer in complex with diploptene. We used the default AF3 parameters for all predictions.

## DISCUSSION

Gene annotation relies on the detection of conserved sequence patterns, typically using probabilistic models such as profile HMMs. The first protein databases relied on manually curated seed sequences to ensure model fidelity, but most of the >30,000 bacterial protein profile HMMs now available in InterPro and other databases were generated without strict oversight and with limited experimental findings. Our benchmarking of profile HMMs for hopanoid biosynthesis enzymes demonstrates that the accuracy of such models is variable and difficult to predict. This finding allowed us to establish a higher fidelity annotation pipeline, and it also provides a cautionary note on the limitations of machine learning that as applications of generative artificial intelligence become more common. An annotation pipeline is necessarily limited by the breadth and quality of the experimental data on which it is trained, and such pipelines must be continuously re-evaluated as new experimental data become available.

Our benchmarked annotation approach allowed us to refine the understanding of the distribution of hopanoid biosynthesis in bacteria. The major revision relative to prior studies on hopanoid distribution^14,44–46^ is that we observe minimal SHC conservation within bacillota. The only SHC-producing species we identified in this phylum are members of the alicyclobacillaceae family such as *Alicyclobacillus acidocaldarius*, which is the only experimentally verified hopanoid-producing firmicute^47,48^. The SHC-like enzymes from bacillota that were classified as SHC in previous works appear to be sporulenol cyclases (sqhC) that do not produce hopanoids; these squalene cyclase families appear to have evolved early in triterpene cyclase evolution but represent functionally distinct groups^49^. We similarly distinguish hopanoid-producing strains from sterol-producing strains, which is particularly relevant to recent analyses of the planctomycetota. New work has proposed that a novel planctomycete isolate from marine algae, *Crateriforma conspicua* Mal65, produces hopanoids to support oxygen-tolerant symbiotic nitrogen fixation^50^. However, we do not observe SHC in any members of the pirellulales order to which *C. conspicua* belongs and instead observed only terpene cyclases involved in sterol production. Given that sterol production is known to occur and is essential in the model planctomycete *Gemmata obscuriglobus*^51^, our results suggest it is unlikely that *C. conspicua* produces hopanoids.

Our conclusions regarding the distribution of methylated hopanoids largely agree with prior work. For HpnR, our analysis slightly expands the range of 3Me-hopanoid-producing organisms to include members of the *Paraburkholderia* genus; however, 3Me-hopanoid production has never been reported in these species. As there is only one *hpnR* gene validated by knockout or biochemical analysis in the literature, limiting the benchmarking confidence, whether *Paraburkholderia* are true 3Me-hopanoid-producers will await future experimental analysis. Regarding 2Me-hopanoids generated by HpnP, our benchmarked distribution appears similar to that of Hoshino *et al.*^14^ in cyanobacteria; however, comparing these distributions required extensive re-analysis due to the use of different taxon nomenclatures and taxon classification criteria. Though whole-genome classification approaches such as GTDB-Tk are widely accepted as more accurate, the resulting classifications can be substantially different from those supported by bacterial taxonomic authorities such as LPSN. Hundreds of new and re-classifications have been formally proposed based on whole-genome definitions^52^, but these name changes have yet to be widely adopted by researchers. We encourage future hopanoid researchers to prioritize the more accurate whole genome-based taxonomies to facilitate comparison of results across publications.

In addition to understanding their distribution, we analyzed the phylogenetic relationships of hopanoid-related enzymes to predict their origins. These relationships suggested that unmethylated hopanoids, including their C_35_ varieties, originated in an ancestor of marine alphaproteobacteria. This was further supported by our determination that hopanoids are strongly associated with adaptations to high osmolarity environments, methanotrophy, and nitrogen cycling. The link between hopanoids and osmotic stress tolerance is expected given the similarity of hopanoids to eukaryotic sterols, which are known to fortify cell membranes to reduce permeability and resist force-mediated lysis. It is well established that hopanoids similarly condense and thicken *in vitro* lipid bilayers^1,2^, and though the relevance of these studies to living membranes is not completely clear, sensitivity to osmotic stress has been reported in nearly every hopanoid-deficient mutant tested^20,53–56^. We speculate that survival of high salinity environments was a major driver of hopanoid evolution.

The enrichment of single carbon- and nitrogen-metabolism in hopanoid-producing organisms is more difficult to parse. Many of these processes rely on isoprenoid quinones or radical SAM enzymes, and it is possible that their correlation with hopanoids arises from the use of shared substrates rather than a functional relationship. Presence of hopanoids in marine nitrite-oxidizing bacteria (NOB) has been recently emphasized in hopanoid geobiology literature^10–12^, and in our growth assays, SHC is required for efficient nitrite utilization in the terrestrial nitrogen-fixing and denitrifying^57^ alphaproteobacterium *Bradyrhizobium diazoefficiens*. Whether hopanoids participate directly in nitrite utilization will require future experiments. Though previous work identified an enrichment of hopanoids in plant-associated nitrogen-fixing bacteria^9^, subsequent work suggests that hopanoids do not play a direct role in nitrogen fixation^20,58^. Rather, we believe that hopanoids and nitrogen fixation are correlated because they are both common to low oxygen environments, given that hopanoids (unlike sterols) do not require oxygen for their biosynthesis and that oxygen is inhibitory to the nitrogen fixation machinery.

The degree to which unmethylated geo-hopane deposition can be attributed to ancestors of marine alphaproteobacteria will be an exciting future area for hopanoid geobiology research. We are only beginning to appreciate the diversity and significance of non-cyanobacterial marine hopanoid producers. Though such species are dominant in marine animal microbiomes^59–61^, and symbiotic alphaproteobacteria appear to be major contributors to modern marine nitrogen cycling^62^, relatively few complete marine alphaproteobacterial genomes are available. Incorporating our growing understanding of the biology marine alphaproteobacteria, ideally coupled with genetic tools, will aid our knowledge of the role of hopanoids in these origins.

The origins of the 2Me- and 3Me-geo-hopanes are less clear. We could not identify a single phylum of origin for either HpnR or HpnP proteins, though all of the statistically supported potential root positions for both protein families were all within the clade of **Figure 1** containing the pseudomonadota, acidobacteriota, and desulfobacterota. Our HpnP tree is similar to the HpnP tree published by Hoshino and colleagues^14^, as both trees indicates that the earliest branching HpnP sequences are from phyla other than cyanobacteriota or alphaproteobacteria. The other group’s conclusion that 2Me-geo-hopanes dated prior to ∼750 Ma were deposited by cyanobacteria is based on a study on the origin of the rhizobiales order of alphaproteobacteria that proposed that the hopanoid-producing families of alphaproteobacteria did not emerge until 750 Ma or later^15^. In our analyses, we observe high levels of HpnP conservation in the following alphaproteobacterial families: xanthobacteraceae, acetobacteraceae, beijerinckaceae, and reyranellaceae; we note the latter family does not appear to be represented in the Hoshino et al. dataset, perhaps due to differences in HpnP annotation approaches. A new geologic timescale of evolution across all bacteria suggests the earliest-branching of these families is the reyranellaceae, originating between 783-1212 Ma (95% confidence interval)^63^. Though this supports the conclusion that the oldest 2Me-geo-hopanes were more likely deposited by cyanobacteria than alphaproteobacteria, we restrict that conclusion to geo-hopanes dated prior to ∼1200 Ma. Whether cyanobacteria were the first of *any* phyla to produce hopanoids, however, cannot be resolved by molecular dating. The same pan-bacterial evolution timeline indicates that the desulfuromonadia class of the desulfobacterota F phlyum originated 937-2040 Ma^63^. HpnP is conserved highly in this class, and this phylum was the most highly supported root position for HpnP in our phylogenetic analyses.

Beyond implications for geo-hopane interpretation, our work suggests new avenues for hopanoid cell biology. We identify four new uncharacterized but likely hopanoid-related proteins: the previously annotated HpnL and HpnM families and candidate proteins HpnS and HpnT, which are new to this study. Using the latest structure prediction tools, we predict that HpnS, HpnM, and HpnT participate in hopanoid trafficking across proteobacterial cell envelopes, interacting with the inner membrane hopanoid transporter HpnN in a manner analogous to the MlaC-MlaA-MlaBDEF phospholipid transport system. New analysis of *mlaC*-like gene copy numbers across bacteria^64^ supports this notion. Tripathi et al. identified a subset of pseudomonadota that contain multiple copies of *mlaC*-like genes. In these species, which we find to be SHC-containing organisms, one *mlaC*-like copy is adjacent to other *mla* genes, whereas additional *mlaC*-like copies typically are adjacent to hopanoid-associated genes. The bacterial genera containing secondary, *hpnN*-adjacent *mlaC*-like genes in Tripathi et al. largely overlap with the genera in which we’ve identified *hpnM* genes, suggesting that we are referring to the same loci. This opens up future directions for cellular, structural, and biochemical assays to test this hypothetical transport pathway. If our functional predictions are verified, it will be interesting to study how these protein families originated, both relative to the Mla proteins and in the alpha- versus gamma-proteobacteria.

The function of HpnL remains unclear. Its closest relatives are the AglD family of lysylphosphatidylglycerol (lysyl-PG) synthases, which covalently link the carboxyl group of lysine to the terminal hydroxyl in the glycerol moiety of PG^65,66^, increasing the positive charge of the bilayer to improve resistance to cationic antimicrobial peptides. It is possible that HpnL might use a similar chemistry to join amino acids or other carboxylated compounds to the polyol groups of extended hopanoids such as bacteriohopanetetral (BHT), though the structure of BHT is quite distinct from the PG substrate of AglD. Covalent attachment of BHT to the amino acid derivative phenyl-acetic acid (PAA), an auxin-like compound and weak phytohormone, has been reported in actinobacterial *Frankia* spp.^67,68^, suggesting other amino acid-hopanoid conjugates exist. However, the HpnL-associated protein domain (TIGR03476) is found predominantly in pseudomonadota, and to our knowledge PAA-BHT not been found in this phylum. The function of HpnL awaits future genetic characterization.

## ACKNOWLEGEMENTS

This work was supported by NIH grants R00GM126141 and R35GM147015 (to B.J.B) and Carnegie Institution for Science endowment funds. Computational analyses were performed using Caltech’s Resnick High Performance Computing Center and the NIH Biowulf HPC. We thank members of the Belin lab, Dr. Karina Gutiérrez-García (University of Arizona), Dr. Will Ludington (Johns Hopkins University), Dr. Damian Ekiert (Johns Hopkins University), and Dr. Dianne Newman (Caltech) for helpful discussions and feedback. We are grateful to Carnegie Embryology’s IT, front office, and facilities support staff for making our work possible.

The contributions of NIH authors were made as a part of their official duties as NIH federal employees, are in compliance with agency policy requirements, and are considered works of the United States Government. However, the findings and conclusions presented in this paper are those of the authors and do not necessarily reflect the views of the NIH or the US Department of Health and Human Services.

---

**Figure S1.**
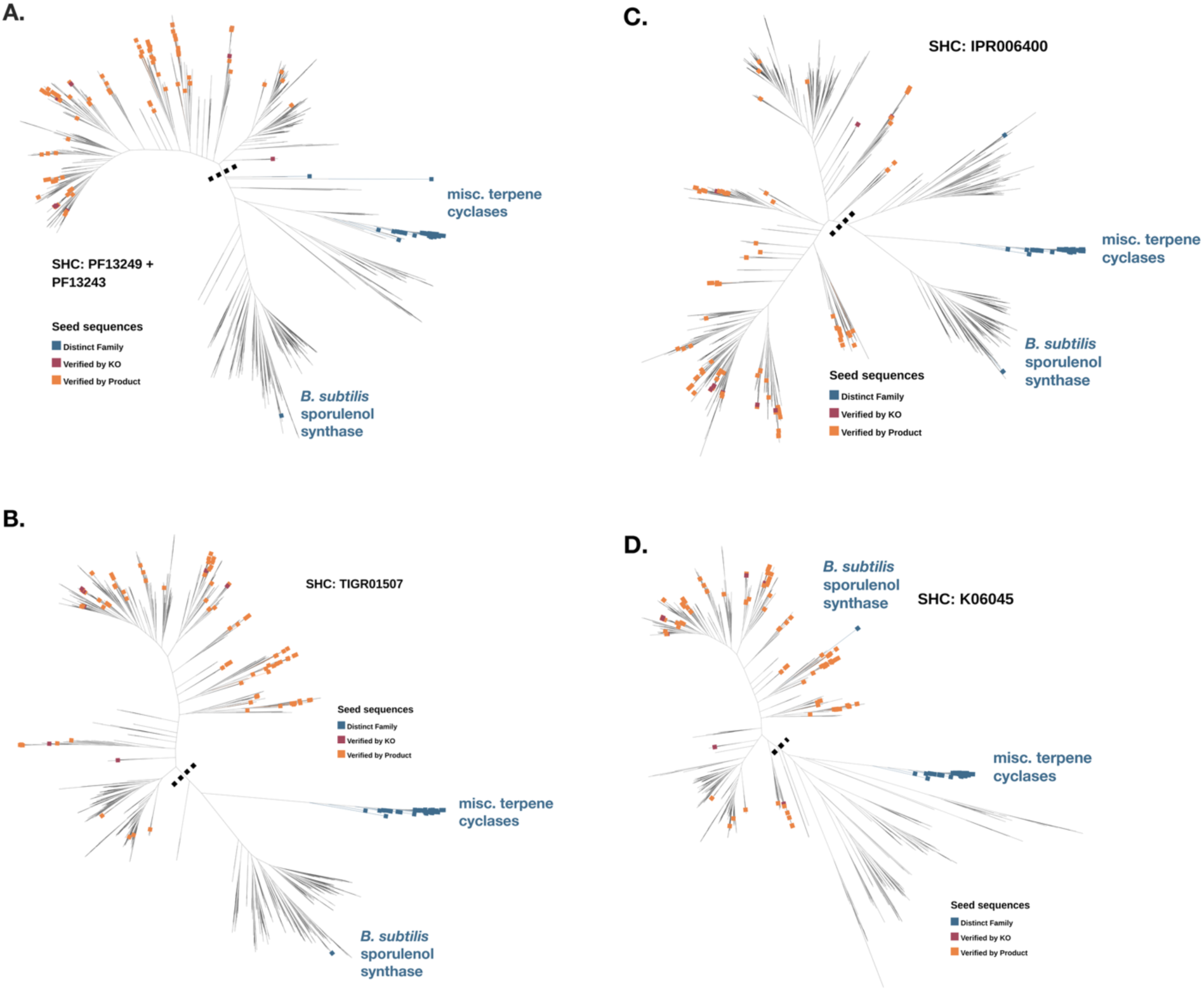
Benchmarking tree for SHC-associated domain profiles. Unrooted maximum likelihood phylogenetic trees of SHC benchmarking control sequences plus proteins associated with PF13249+PF13243 (**A**), TIGR01507 (**B**), IPR006400 (**C**), K06045 (**D**) in UniProt or IMG. Negative control sequences (“Distinct family”, blue), knockout- or biochemically-validated positive control sequences (“Verified by KO”, red), and positive control sequences verified by presence of hopanoid product in culture (“Verified by Product”, orange) are indicated by box labels at branch tips. Dotted black line indicates position of separation between positive and negative control-associated clades.

**Figure S2.**
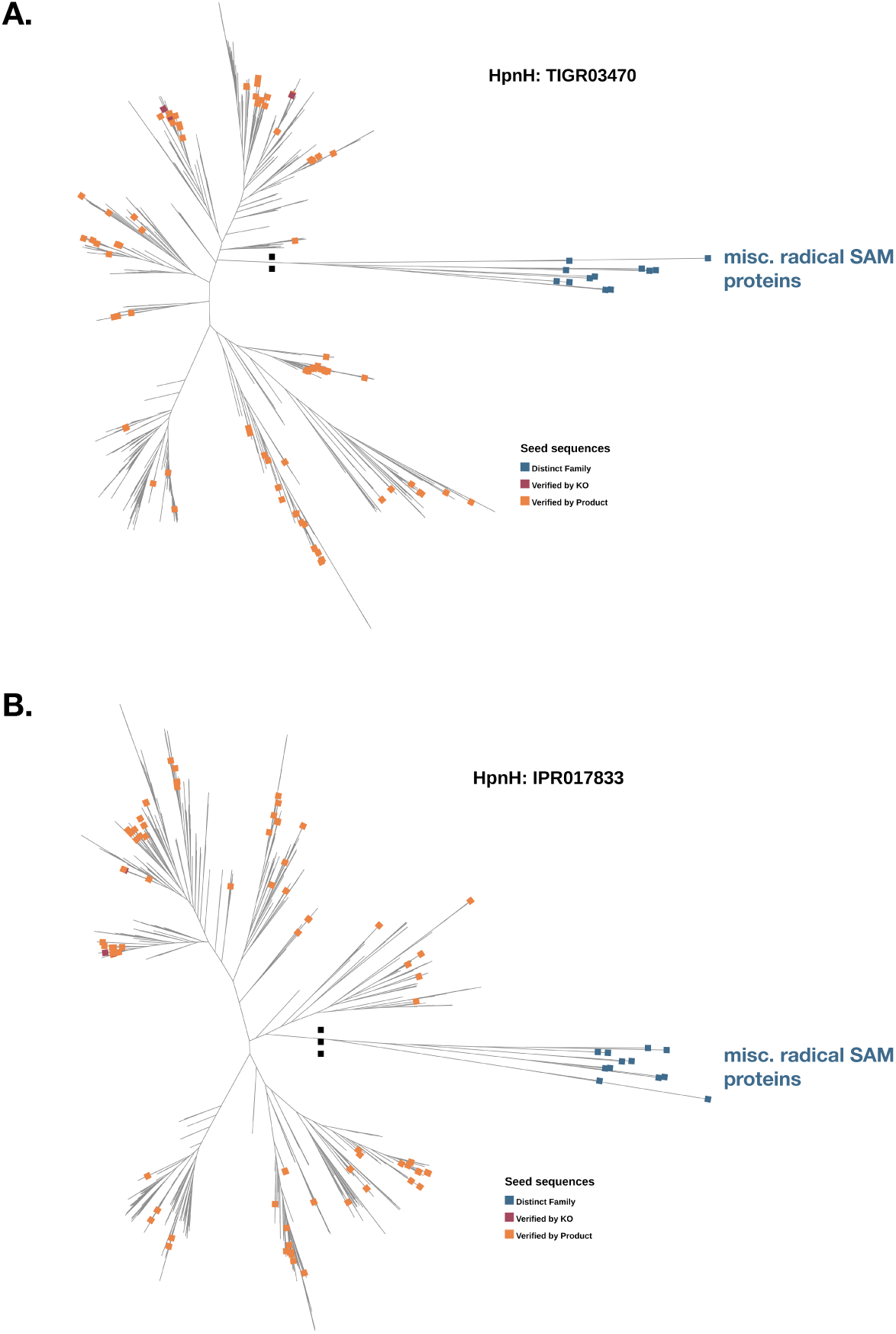
Benchmarking tree for HpnH-associated domain profiles. Unrooted maximum likelihood phylogenetic trees of SHC benchmarking control sequences plus proteins associated with TIGR03470 (**A**) or IPR017833 (**B**) in UniProt or IMG. Negative control sequences (“Distinct family”, blue), knockout- or biochemically-validated positive control sequences (“Verified by KO”, red), and positive control sequences verified by presence of hopanoid product in culture (“Verified by Product”, orange) are indicated by box labels at branch tips. Dotted black line indicates position of separation between positive and negative control-associated clades.

**Figure S3.**
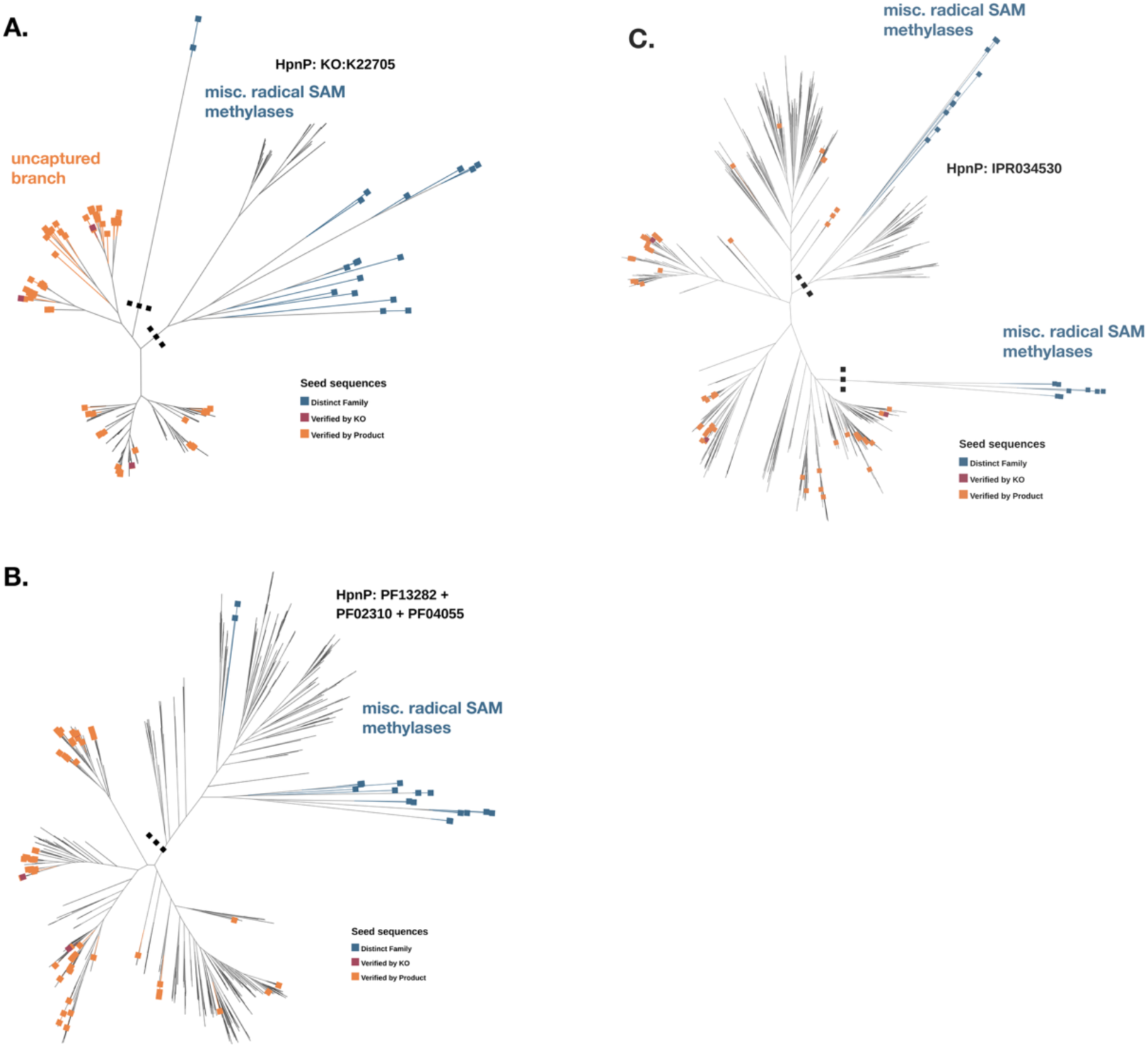
Benchmarking tree for HpnP-associated domain profiles. Unrooted maximum likelihood phylogenetic trees of SHC benchmarking control sequences plus proteins associated with PF13282+PF02310+PF04055 (**A**), IPR034530 (**B**), or K22705 (**C**) in UniProt or IMG. Negative control sequences (“Distinct family”, blue), knockout- or biochemically-validated positive control sequences (“Verified by KO”, red), and positive control sequences verified by presence of hopanoid product in culture (“Verified by Product”, orange) are indicated by box labels at branch tips. Dotted black line indicates position of separation between positive and negative control-associated clades.

**Figure S4.**
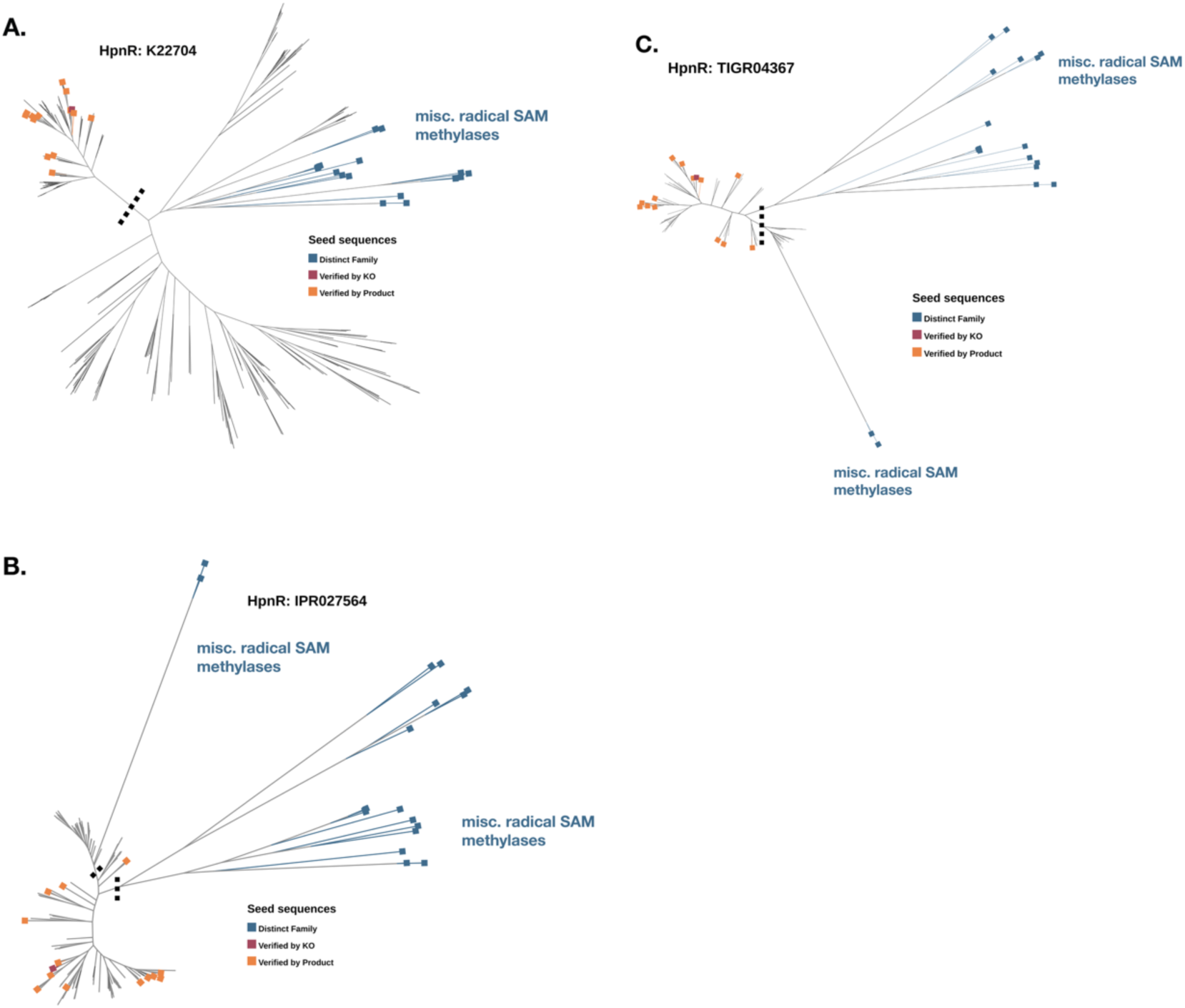
Benchmarking tree for HpnR-associated domain profiles. Unrooted maximum likelihood phylogenetic trees of SHC benchmarking control sequences plus proteins associated with K22704 (**A**), IPR027564 (**B**), or TIGR04367 (**C**) in UniProt or IMG. Negative control sequences (“Distinct family”, blue), knockout- or biochemically-validated positive control sequences (“Verified by KO”, red), and positive control sequences verified by presence of hopanoid product in culture (“Verified by Product”, orange) are indicated by box labels at branch tips. Dotted black line indicates position of separation between positive and negative control-associated clades.

**Figure S5.**
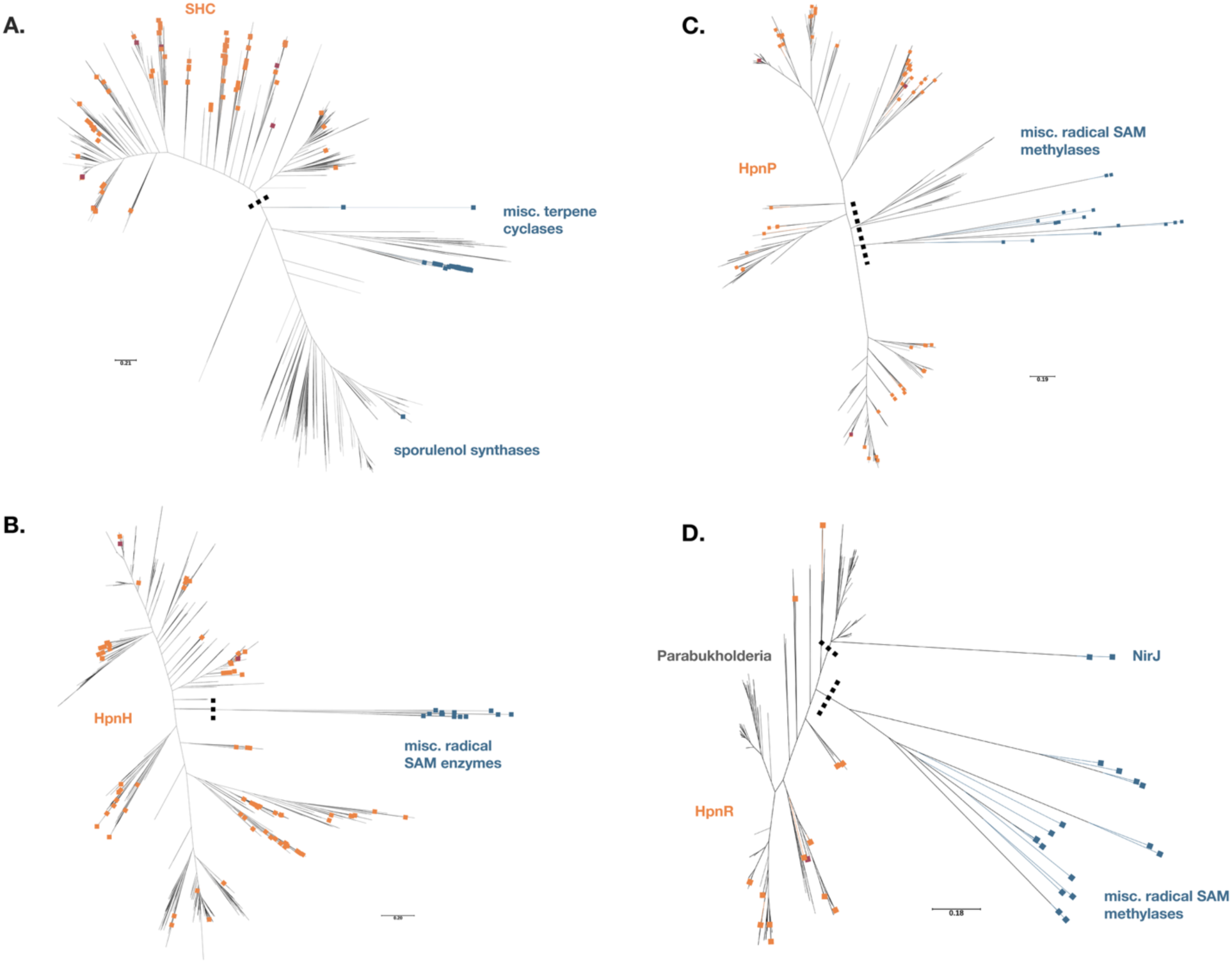
Benchmarking tree for identifying SHC, HpnH, HpnP, and HpnR family members in IMG genomes. Unrooted maximum likelihood phylogenetic trees of benchmarking control sequences plus proteins associated with (**A**) SHC (Pfam13249+Pfam13243), (**B**) HpnH (TIGR03470), (**C**) HpnP (IPR034530), and (**D**) HpnR (TIGR04367) in genomes from Table S3. Negative control sequences (“Distinct family”, blue), knockout-or biochemically-validated positive control sequences (“Verified by KO”, red), and positive control sequences verified by presence of hopanoid product in culture (“Verified by Product”, orange) are indicated by box labels at branch tips. Dotted black line indicates position of separation between positive and negative control-associated clades. Scale bars indicate substitutions per 1000 amino acids.

**Figure S6.**
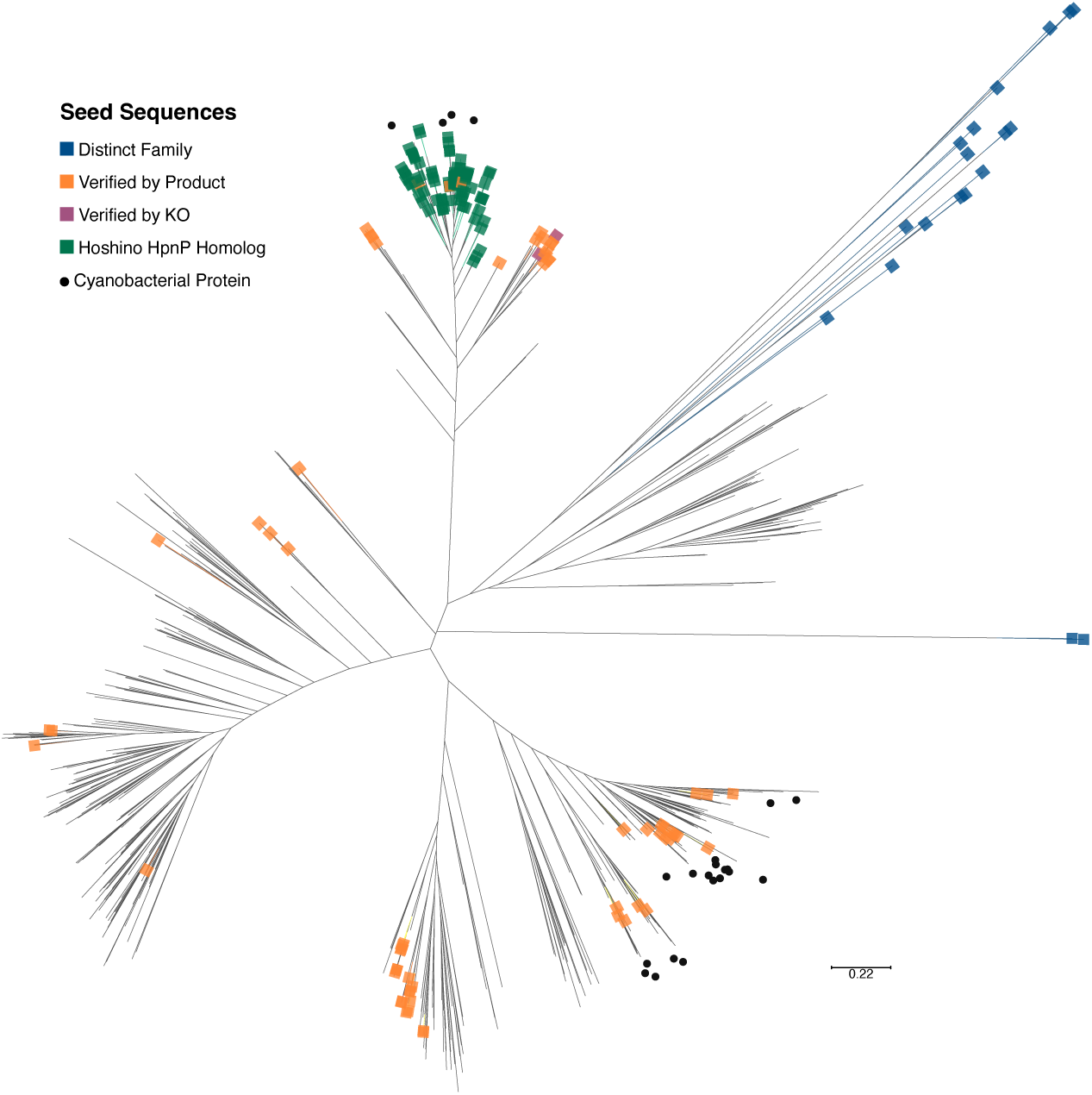
Comparison of IMG HpnP domain-associated proteins with Hoshino *et al.* HpnP homologs. Unrooted maximum likelihood phylogenetic tree of HpnP benchmarking control sequences, IPR034530-annotated proteins in IMG, and cyanobacterial HpnP homologs (annotated with green rectangles at branch tips) identified by Hoshino *et al*. (annotated with green rectangles at branch tips) ^14^. Control sequences are annotated as in Fig. S1-S5. Experimentally validated HpnP proteins in the control sequences are indicated by black circles, demonstrating that cyanobacterial HpnP proteins form two clades; only one clade is captured by the Hoshino *et al*. annotation method.

**Figure S7.**
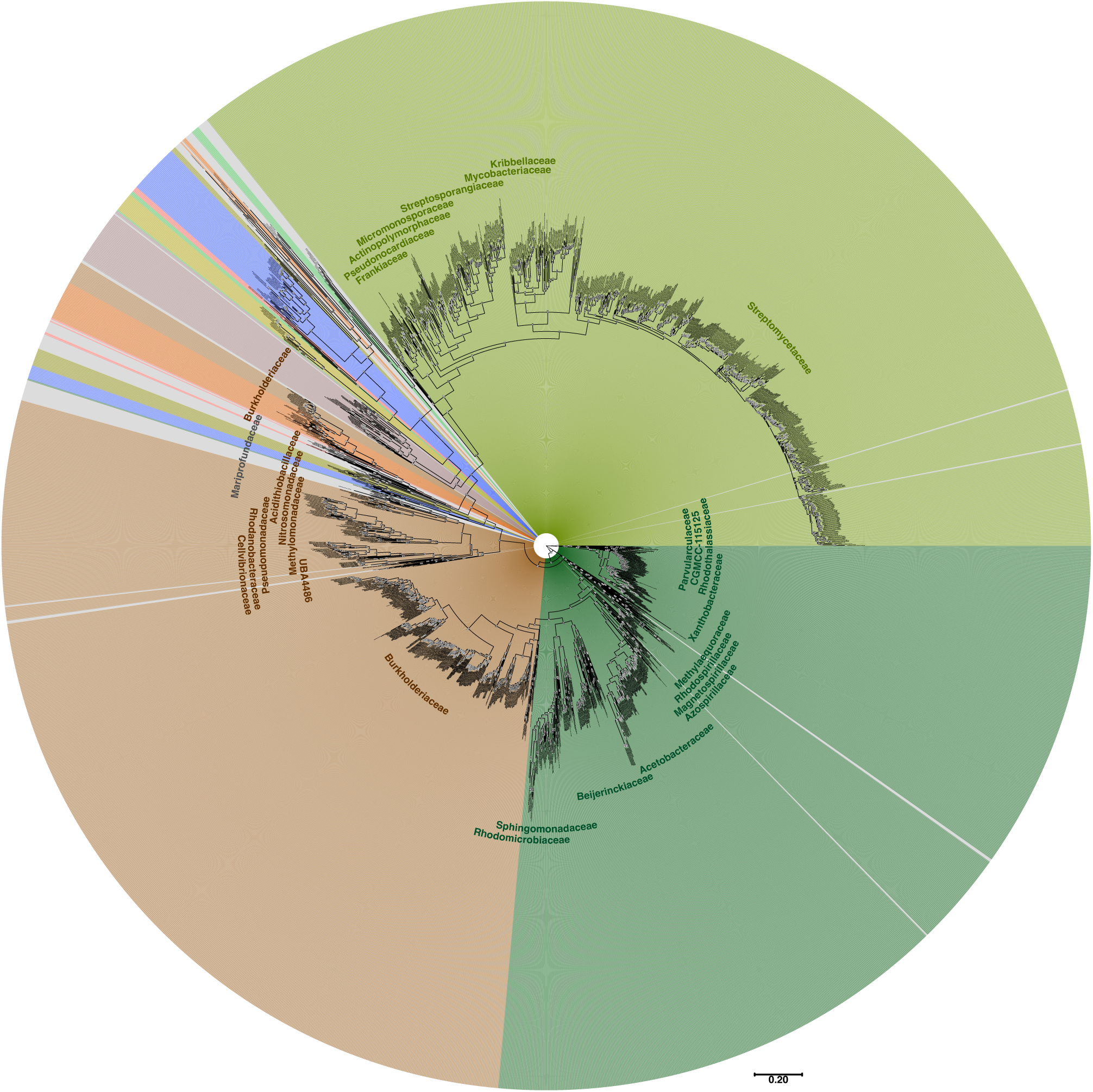
Extended hopanoids originated in marine alphaproteobacteria. Maximum likelihood phylogenetic tree generated for protein sequences associated with HpnH. Branch names include IMG genome ID, GTDB family and genus assignments, and IMG protein ID. Numbers at internal nodes (highlighted in white) indicate branch support values using ultrafast bootstrap approximation (UFBoot), based on 1000 bootstraps; 100 = highest confidence, 0 = no confidence. Scale bar indicates substitutions per 1000 amino acids. Branch backgrounds are color-coded by phyla according to the color scheme (outer ring and phylum text labels) used in Figure 1.

**Figure S8.**
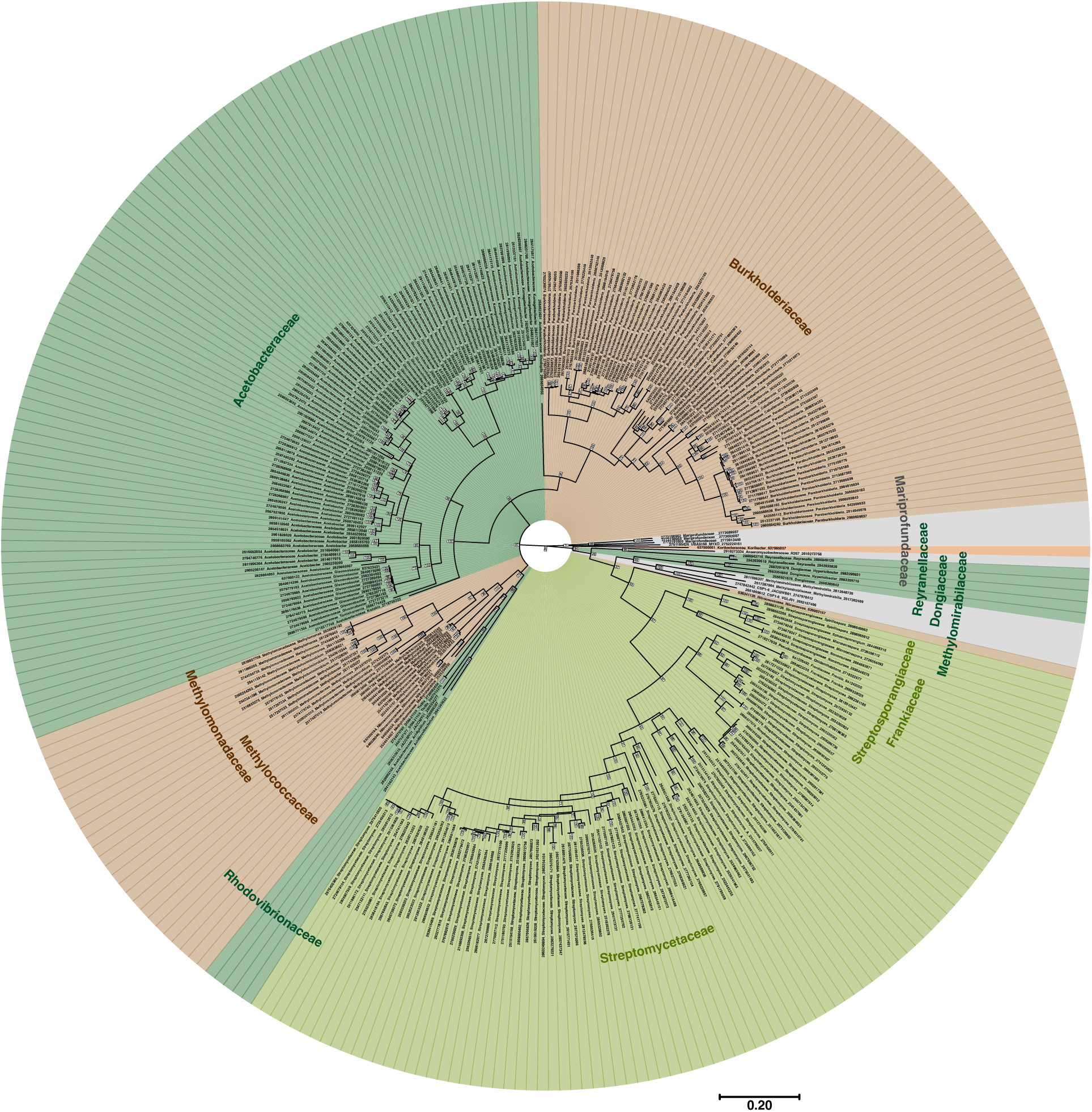
Phylogenetic relationships of HpnR homologs. Maximum likelihood phylogenetic tree generated for protein sequences associated with HpnR. Branch names include IMG genome ID, GTDB family and genus assignments, and IMG protein ID. Numbers at internal nodes (highlighted in white) indicate branch support values using ultrafast bootstrap approximation (UFBoot), based on 1000 bootstraps; 100 = highest confidence, 0 = no confidence. Scale bar indicates substitutions per 1000 amino acids. Branch backgrounds are color-coded by phyla according to the color scheme (outer ring and phylum text labels) used in Figure 1.

**Figure S9.**
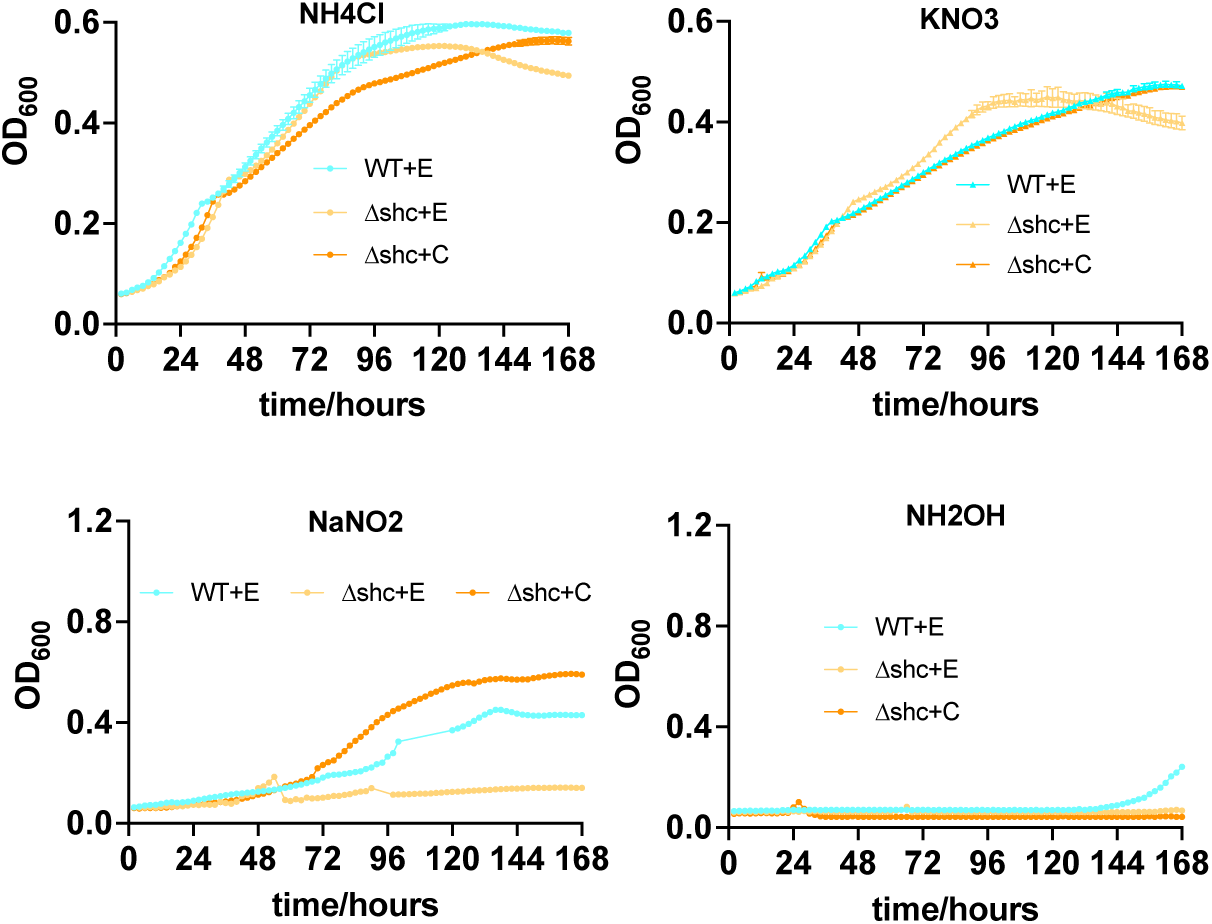
Hopanoid-deficient *B. diazoefficiens* have slow growth on nitrite. Growth curves of WT (WT+E), hopanoid-deficient (delta *shc* + E), and rescue (delta *shc* + C) strains of *B. diazoefficiens* on minimal media supplemented with indicated nitrogen sources.

**Figure S10.**
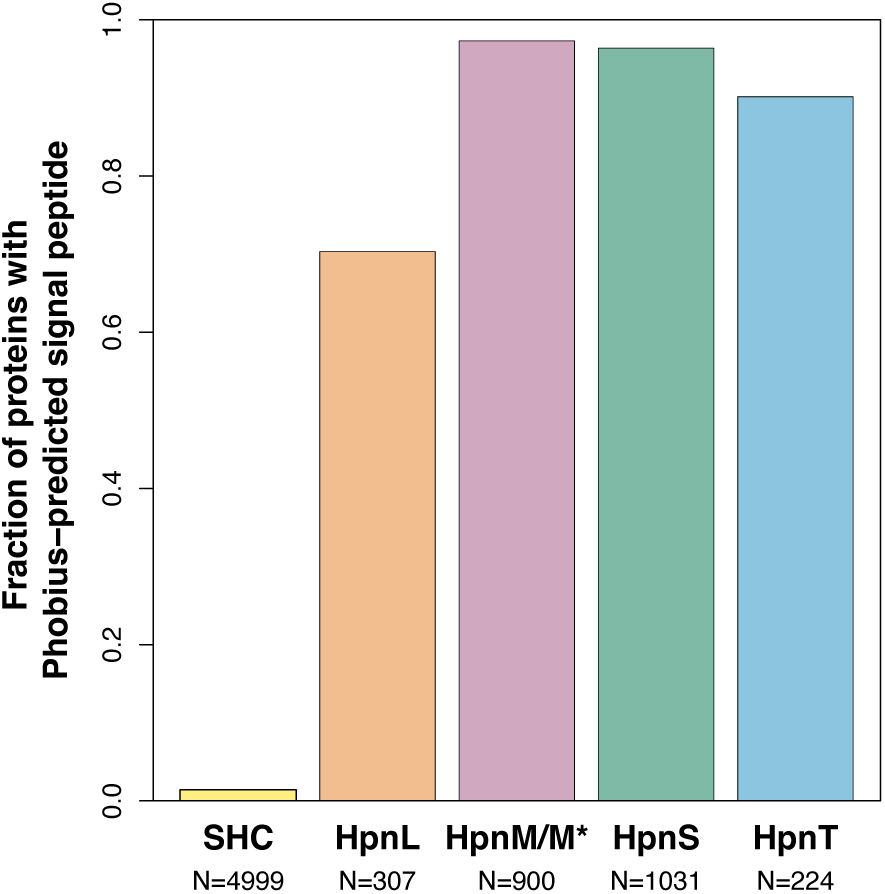
Signal peptides are prevalent in HpnL, HpnM, HpnS, and HpnT proteins. Fraction of proteins each family predicted to have a signal peptide for periplasmic secretion by Phobius^78^.

## Notes

### Competing Interest Statement

The authors have declared no competing interest.

### Summary of Updates

New analyses performed; Figures 1-3 revised; supplemental files added; supplemental files updated.

